# Single-cell image-based genetic screens systematically identify regulators of Ebola virus subcellular infection dynamics

**DOI:** 10.1101/2024.04.06.588168

**Authors:** Rebecca J. Carlson, J.J. Patten, George Stefanakis, Brian Y. Soong, Adityanarayanan Radhakrishnan, Avtar Singh, Naveen Thakur, Gaya K. Amarasinghe, Nir Hacohen, Christopher F. Basler, Daisy Leung, Caroline Uhler, Robert A. Davey, Paul C. Blainey

**Affiliations:** Massachusetts Institute of Technology, Department of Health Sciences and Technology, Cambridge, MA, USA; Broad Institute of MIT and Harvard, Cambridge, MA, USA; Department of Virology, Immunology, and Microbiology, Boston University School of Medicine, Boston, MA, USA; National Emerging Infectious Diseases Laboratories, Boston University, Boston, MA, USA; Laboratory for Information & Decision Systems, Massachusetts Institute of Technology, Cambridge, MA, USA; Harvard School of Engineering and Applied Sciences, Cambridge, MA, USA; Department of Microbiology, Icahn School of Medicine at Mount Sinai, New York, NY, USA; Division of Infectious Diseases, Washington University School of Medicine, St. Louis, MO, USA; Massachusetts General Hospital, Cancer Center, Boston, MA, USA; Massachusetts Institute of Technology, Department of Biological Engineering, Cambridge, MA, USA; Koch Institute for Integrative Cancer Research, MIT, Cambridge, MA, USA

## Abstract

Ebola virus (EBOV) is a high-consequence filovirus that gives rise to frequent epidemics with high case fatality rates and few therapeutic options. Here, we applied image-based screening of a genome-wide CRISPR library to systematically identify host cell regulators of Ebola virus infection in 39,085,093 million single cells. Measuring viral RNA and protein levels together with their localization in cells identified over 998 related host factors and provided detailed information about the role of each gene across the virus replication cycle. We trained a deep learning model on single-cell images to associate each host factor with predicted replication steps, and confirmed the predicted relationship for select host factors. Among the findings, we showed that the mitochondrial complex III subunit UQCRB is a post-entry regulator of Ebola virus RNA replication, and demonstrated that UQCRB inhibition with a small molecule reduced overall Ebola virus infection with an IC50 of 5 μM. Using a random forest model, we also identified perturbations that reduced infection by disrupting the equilibrium between viral RNA and protein. One such protein, STRAP, is a spliceosome-associated factor that was found to be closely associated with VP35, a viral protein required for RNA processing. Loss of STRAP expression resulted in a reduction in full-length viral genome production and subsequent production of non-infectious virus particles. Overall, the data produced in this genome-wide high-content single-cell screen and secondary screens in additional cell lines and related filoviruses (MARV and SUDV) revealed new insights about the role of host factors in virus replication and potential new targets for therapeutic intervention.

## Introduction

Ebolaviruses such as Zaire Ebola virus (EBOV), Sudan virus (SUDV), and the distantly related Marburg virus (MARV), are single-stranded, negative-sense filoviruses responsible for outbreaks with high case fatality rates, predominantly in West or equatorial Africa (Ilunga Kalenga et al., 2019; Lo et al., 2017). EBOV infection results in Ebola virus disease (EVD), characterized by inflammatory responses, immunosuppression, and major fluid losses (Malvy et al., 2019). Monoclonal antibody therapy has demonstrated modest efficacy, reducing EVD case fatality rates to 30% (Mulangu et al., 2019). While a recently FDA-approved live-attenuated recombinant VSV vaccine expressing EBOV glycoprotein (GP) resulted in EBOV-specific protection (Heppner et al., 2017; Mulangu et al., 2019), there are no approved vaccines against MARV or SUDV.

Genetic screens are high-throughput methods that enable identification of host targets that modulate viral infection. However, due to the challenges of screening high-consequence viruses, previous genetic screens for Ebola virus often relied on the use of pseudotyped viruses (Bruchez et al., 2020; Carette et al., 2011; Cheng et al., 2015), precluding identification of post-entry viral modulators, or on model virus genomes and reporter expression (Martin et al., 2018), which may not fully recapitulate the live Ebola virus life cycle. Two genome-wide screens that used live Ebola virus (Filone et al., 2015; Flint et al., 2019) both assessed virus-induced cell death, which for Ebola virus takes many days, and relied on selection and growth of surviving cells, a complex outcome biased towards recovery of near-complete blocks to virus replication seen upon loss of cell entry factors. Here, we set out to assess cell survival-independent phenotypes, allowing for recovery of host factors affecting any part of the virus replication cycle and enabling discovery of a larger range of therapeutic intervention points.

Our optical pooled screening (OPS) (Feldman et al., 2019) approach enables image-based pooled genetic screens in tens of millions of cells (Carlson et al., 2023; Funk et al., 2022). OPS couples high-resolution images of single cells paired with targeted *in situ* sequencing readout of each cell’s specific genetic perturbation identity. In the pooled perturbation format, this approach scales to large numbers of cells and perturbations. This high throughput enables statistically high-powered analyses of the role of many individual host factors in cellular phenotypes at the cellular or subcellular levels. The high-content data produced by such image-based genetic screens can be used directly to derive meaningful insights into genetic function. Such insights can further support well-informed prioritization of hits for productive allocation of resources in downstream mechanistic studies (Carlson et al., 2023; Funk et al., 2022). Here, we present the first genome-wide multiparametric genetic screen for EBOV, evaluating viral protein and RNA synthesis as markers of infection, whose results characterize an extensive landscape of the effects of hundreds of host genes on EBOV replication. We apply machine learning approaches, including deep learning, to our image-based dataset of nearly 40 million single cells and identify regulators of distinct stages of the EBOV lifecycle, from cell entry, to inclusion body formation, and viral RNA transcription and replication. We confirmed hits via secondary screening in two cell lines with EBOV and across two distantly related filoviruses (SUDV and MARV) to robustly validate, generalize, and contextualize our results. UQCRB and STRAP, proteins respectively involved in the mitochondrial respiratory chain and the spliceosome, were selected for further mechanistic analysis of their roles in virus replication.

## Results

### A genome-wide image-based genetic screen reveals regulators of distinct steps in the EBOV replication cycle

EBOV is taken up into cells via macropinocytosis and then trafficked through endolysosomes. Following acidification, host cell cathepsins cleave the viral glycoprotein (GP) into an active form that binds to the intracellular host receptor NPC1, resulting in membrane fusion and cytoplasmic release of the viral capsid (Hoenen et al., 2019). Transcription from the negative stranded virus genome produces viral mRNAs and, in turn, production of viral proteins leads to formation of inclusion bodies (IBs), cytoplasmic foci that serve as sites for viral RNA synthesis (Hoenen et al., 2012, 2019; Nanbo et al., 2013). The EBOV nucleocapsid protein (NP) induces formation of these IBs (Miyake et al., 2020; Wu et al., 2023) and the viral polymerase cofactor VP35 interacts with NP to regulate IB formation (Leung et al., 2015; Miyake et al., 2020). At later stages of infection, these proteins exhibit a diffuse cytoplasmic localization pattern and, finally, localize to the cell periphery during virus budding (Nanbo et al., 2013). These distinct stages of infection are not readily distinguishable by the use of simple intensity-based measurements, for example those typically employed in pooled flow cytometry-based screens. Genome-scale arrayed image-based screens suffer from batch effects that reduce sensitivity to true effects, and are further prohibitively costly and labor-intensive in most research laboratories.

Here we applied OPS to define the contributions of host factors to distinct steps of the EBOV replication cycle at the genome-wide scale, obtaining images of nearly 40 million individual EBOV-infected HeLa-TetR-Cas9 cells transduced with a custom pool of ∼80,000 sgRNAs targeting ∼20,000 genes, including 454 non-targeting control sgRNAs (**Fig. 1a**). Cells were infected with wild-type EBOV in a maximum biocontainment laboratory; subsequently, we assayed the expression level and patterns of EBOV VP35 protein using immunofluorescence (IF) with a monoclonal antibody developed against recombinant VP35 and the Ebola VP35 positive-sense RNA, which predominantly represents EBOV mRNA transcripts (Galão et al., 2022), using fluorescence *in situ* hybridization (FISH) (**Fig. S1a**). EBOV infections were optimized to obtain >90% infection rates as evidenced by VP35 RNA and protein expression (**Fig. S1b**). In addition to measuring viral protein and RNA, we stained for the host transcription factor c-Jun, whose activity is increased in EBOV-and MARV-infected cells (Hölzer et al., 2016; Wynne et al., 2017), as well as LAMP1, a lysosomal protein, vimentin, which is useful for cell segmentation, and a nuclear stain (DAPI) (**Fig. 1b**). Since we had not previously incorporated FISH-based staining in OPS, FISH was optimized as part of our integrated IF and *in situ* sequencing workflow (**Fig. S1a**). To do so, we hybridized primary probes to viral RNA transcripts prior to targeted reverse transcription of sgRNAs. We then amplified the viral RNA signal via hybridization chain reaction (HCR) targeted to the primary probes as previously described (Choi et al., 2018), but with the omission of dextran sulfate, which inhibits the reverse transcriptase and polymerase activities necessary for readout of sgRNA sequences by *in situ* sequencing (Viennois et al., 2013). As expected, cells receiving sgRNAs targeting the Ebola virus receptor NPC1 demonstrated robust loss of both the VP35 RNA and protein signals, and low levels of c-Jun in the cell nucleus (**Fig. 1c, d**).

**Figure 1.**
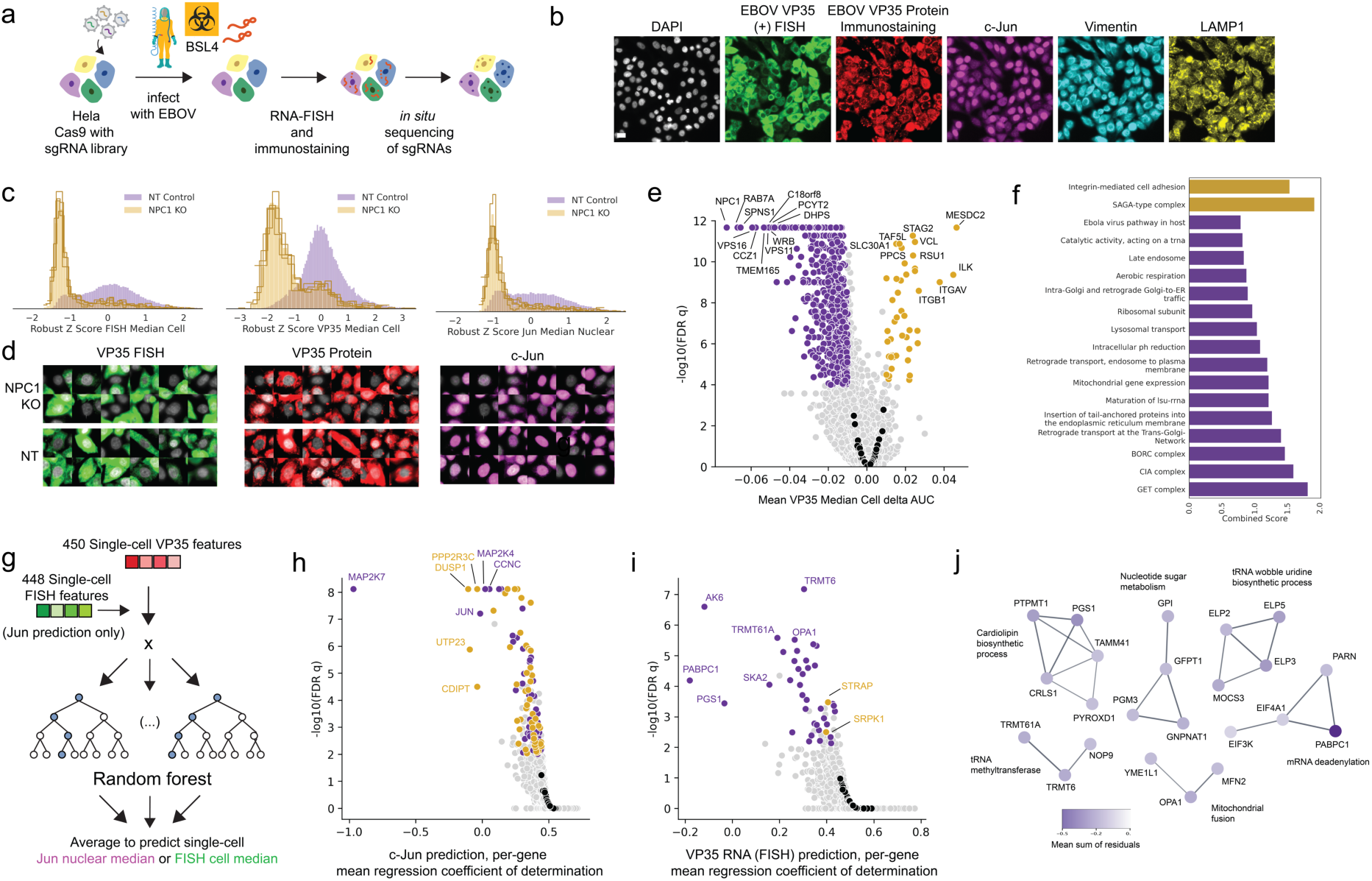
Genome-wide optical pooled screening reveals regulators of multiple responses to Ebola virus infection. (A) Workflow for genome-wide optical pooled screen. (B) Example images of infected cells assayed across six distinct markers in the screen. Scale bar 20 μm. (C) Histograms of VP35 RNA FISH, VP35 protein, and c-Jun transcription factor intensity levels in non-targeting or NPC1 KO cells from the screen, each histogram trace represents a distinct sgRNA targeting NPC1. (D) Randomly selected cells transduced with a non-targeting sgRNA or NPC1-targeting sgRNA show reduced levels of VP35 RNA (FISH), VP35 protein, and c-Jun in NPC1 knockout cells relative to non-targeting cells. (E) Volcano plot of the per-cell median VP35 protein intensity delta AUC between each gene and non-targeting control cells, black points represent distinct non-targeting control sgRNAs. (F) Enrichr gene ontology analysis of top terms significantly enriched in genes that showed reduced VP35 protein intensity upon knockout (purple) or increased VP35 protein intensity (gold). (G) Workflow for random forest regression model trained to predict either c-Jun nuclear intensity from VP35 RNA and protein features or VP35 RNA intensity from VP35 protein features. (H) Volcano plot of random forest regression model coefficients of determination for the c-Jun prediction task, black points represent distinct non-targeting control sgRNAs; genes in purple had a negative mean sum of residuals, indicating decreased c-Jun relative to model prediction, while genes in gold had a positive mean sum of residuals. (I) Volcano plot of random forest regression model coefficients of determination for the VP35 RNA prediction task, black points represent distinct non-targeting control sgRNAs. (J) STRING analysis of genes that had a negative mean sum of residuals for the VP35 RNA FISH prediction task; purple shade denotes magnitude of mean sum of residuals, indicating the amount that EBOV VP35 positive-sense RNA was decreased relative to protein levels. Edge thickness corresponds to confidence score; only interactions with a confidence score >= 0.7 in the full STRING v11.5 were considered.

We identified 998 genetic knockouts that significantly impact Ebola virus infection (FDR-adjusted p-value < 1e-4) as measured by a VP35 protein intensity statistic based on the difference in cumulative area under the curve (delta AUC), as previously described (Feldman et al., 2019). The delta AUC between the cumulative distribution of per-cell VP35 median intensity for each sgRNA relative to non-targeting control cells is a robust metric that captures shifts in the distributions of single-cell intensity values (**Fig. 1e**, **Table S1**, **Table S2**). Aside from NPC1, other genes previously shown to be required for EBOV infection were identified as top hits, including all six members of the HOPS complex, which regulates endolysosomal vesicle fusion (all six scored in the top 60 genes). Only one previous EBOV screen recovered all six members (Carette et al., 2011), highlighting the robustness of our screen and its sensitivity in the genome-wide setting. CTSB and CTSL, required for GP cleavage prior to NPC1 binding, and other previously identified EBOV entry regulators (SPNS1, GNPTAB, UVRAG, PIKFYVE, FIG4, and EXT1) also scored (Carette et al., 2011; Cheng et al., 2015; Filone et al., 2015; Flint et al., 2019), in addition to CAD, an enzyme critical for pyrimidine biosynthesis that was previously identified only in a screen using a synthetic genome replication system (Martin et al., 2018).

In addition to these factors, many genes and protein complexes not previously reported to regulate EBOV infection scored in our analysis of cellular VP35 protein expression levels (**Fig. S1c**) including PIK3C3, involved in autophagy and membrane trafficking; TIMM10, a member of the mitochondrial inner membrane translocase previously shown to interact with VP40 (Batra et al., 2018); as well as entire protein complexes including the conserved oligomeric golgi complex (COG1-8), the GET complex (GET1-4), the GARP complex (VPS51-54), retromer (VPS26A, 29, 35; previously shown to be required for pseudotyped EBOV entry) (Poston et al., 2022), the CIA complex (MMS19 and CIAO2B), genes involved in heparan sulfate synthesis (NDST and UGDH, likely required for virus attachment to cells (O’Hearn et al., 2015)), and poorly characterized proteins such as TM9SF2 and PTAR1 (**Fig. 1f**). Our VP35 protein intensity-based metric also enabled identification of 57 negative regulators, none of which were previously shown to affect EBOV infection. These novel negative regulators were enriched for chaperones including HSP90B1, MESDC2, and UNC45A (UNC45A was previously shown to interact with Ebola VP30 (J. Fang et al., 2022)), integrin-related genes (such as ITGB1, ITGB5, and ITGAV), mRNA deadenylases (CNOT10, 11), and SAGA complex members (TADA2, TAF5L, TAF6L, TADA2B, SUPT20H) which modulate replication of other RNA viruses (Carlson et al., 2023; Guo et al., 2020) (**Fig. 1f**).

Given that we produced statistically high-powered datasets identifying many regulators of VP35 protein levels as well as VP35 RNA and c-Jun levels for each cell, we next sought to identify genes that differentially regulated VP35 RNA or c-Jun nuclear translocation relative to VP35 protein levels, reasoning that such a dependency analysis may highlight genes with primary effects on cellular responses to infection rather than on viral replication itself. We specified features from the VP35 RNA and protein color channels of the single-cell images using CellProfiler-derived intensity, colocalization, size, shape, and subcellular distribution image features (Bray et al., 2016) as previously described (Funk et al., 2022). We then trained two separate random forest regression models to predict the c-Jun nuclear intensity from the VP35 RNA and protein features (**Fig. 1g, h**) as well as the VP35 RNA intensity from the VP35 protein features (**Fig. 1i**, **Table S3**). By looking for factors that, when targeted, resulted in uncoupling of these features that are typically correlated during infection, we systematically identified genes that separately regulated these responses. As expected, factors where the c-Jun protein levels were poorly predicted by viral VP35 and RNA levels included Jun itself as well as known MAP kinase pathway members (**Fig. 1h**). The gene with the strongest reduction in VP35 RNA relative to VP35 protein levels was the mRNA binding protein PABPC1, which was shown to interact with recombinantly expressed EBOV NP (García-Dorival et al., 2016) and was downregulated in EBOV-infected NHP monocytes (Kotliar et al., 2020). It is unclear if this interaction is through an RNA intermediate, to which each is known to bind. A number of other genes with functions related by STRING analysis also decreased EBOV VP35 RNA relative to protein levels (**Fig. 1j**), while only two genes, SRPK1 and STRAP, had the opposite effect, showing elevated RNA relative to protein (**Fig. 1i**). Notably, SRPK1 was previously shown to regulate EBOV transcription through phosphorylation of VP30 (Takamatsu et al., 2020).

### Deep Neural Network Model Reveals Regulators of Ebola Virus Subcellular Protein Localization

While overall VP35 protein and RNA levels were informative, the single-cell images contain more information about virus replication, including, for instance, subcellular localization patterns of these and additional markers as well as colocalization information. Others have shown that such patterns change over time and likely correspond to remodeling organelles for production of virions and altering the cell to disarm innate antiviral immune responses (Nanbo et al., 2013). Due to the complexity of these localization patterns, we sought to identify regulators of EBOV infection dynamics using a deep learning model. Autoencoders are classical deep learning models used to generate informative representations of the data in an unsupervised fashion (Baldi, 2012). We used a convolutional autoencoder consisting of 5 encoding and decoding layers and a 2048-dimensional bottleneck. The autoencoder was trained to reconstruct all six-channel masked images of single cells. This corresponds to the unsupervised model in **Fig. 2a** and **Fig. S2a-b**. To obtain more informative embeddings, we fine-tuned this model with a supervised objective function to predict four human-labeled categories of EBOV VP35 protein subcellular localization, namely 1) faint, indicative of uninfected cells, 2) punctate, which represents cells with viral inclusion bodies at early stages of infection, 3) cytoplasmic, representing a later stage of infection with diffuse protein localization, and 4) peripheral, representing viral budding. These were manually annotated for over 3,000 cells for model training. PHATE (Moon et al., 2019) was then used to visualize the 2048-dimensional embedding of cells obtained from the two deep learning models: the entirely unsupervised autoencoder model as well as the autoencoder model which was subsequently fine-tuned using a supervised objective. While the unsupervised cell embeddings showed a lack of separation of the four phenotypic classes, the fine-tuned embeddings using the supervised objective showed clear separation among the four classes (**Fig. 2b**). An ordinal chi square test was used to identify genes that significantly altered the proportion of cells in each of the four phenotypic classes representing earlier or later stages of infection relative to non-targeting controls (**Fig. 2c**, **Table S4**).

**Figure 2.**
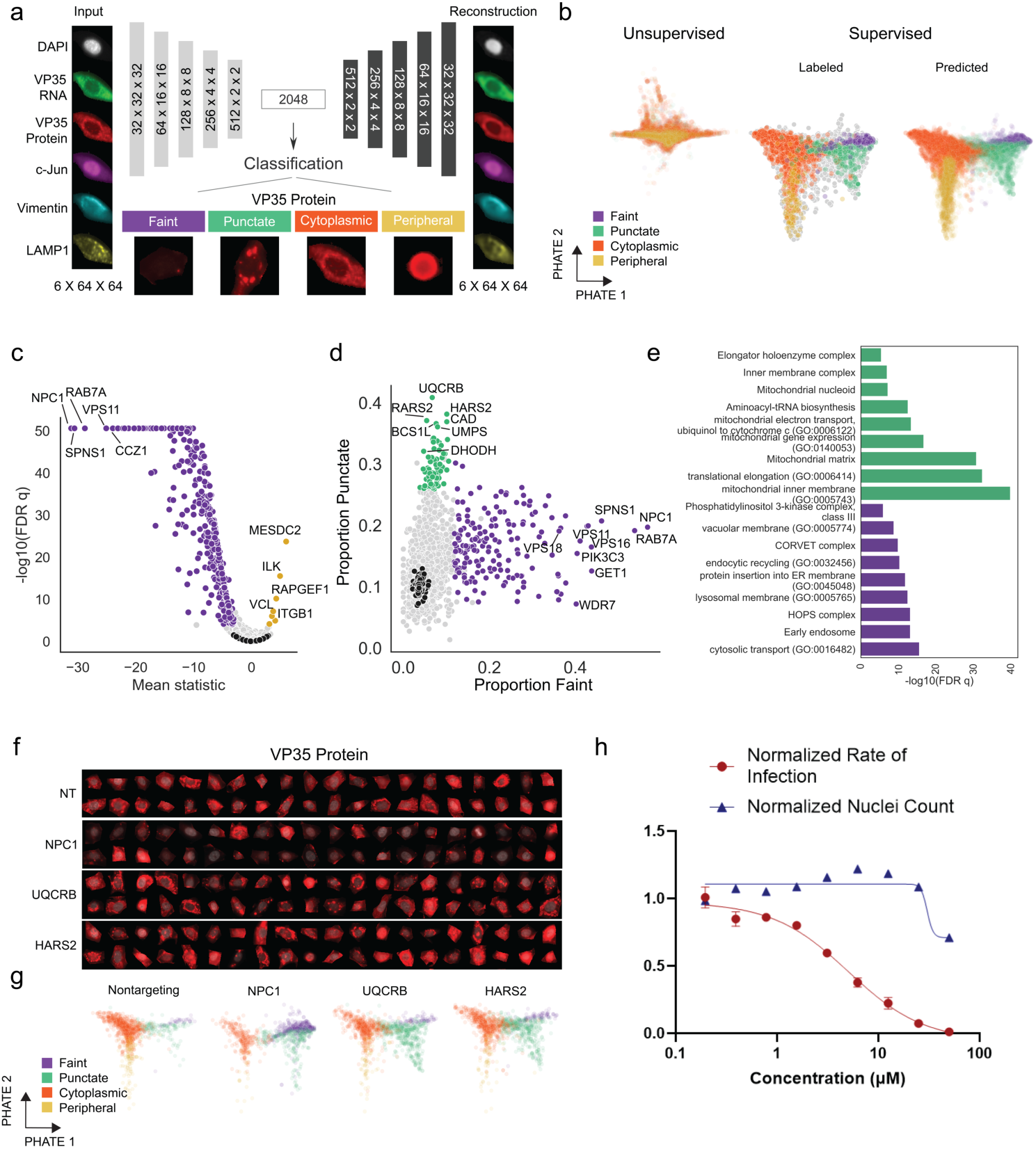
A deep neural network model reveals regulators of Ebola virus VP35 protein subcellular localization. (A) Architecture of the neural network model, trained first as an autoencoder with a latent space of 2048 dimensions to reconstruct input cell images (example input cell and autoencoder reconstruction shown) and subsequently fine-tuned to classify cells based on the manually annotated four classes of VP35 protein localization, with one example image shown for each class. (B) PHATE visualization of the single cell autoencoder embedding (2048 dimensions/cell) obtained from the two deep learning modes: fully unsupervised autoencoder model (left) colored by the predicted class labels, and fine-tuned autoencoder model using the supervised objective function (right) showing both the hand-labeled training data as well as the predicted labels. (C) Volcano plot of per-gene mean ordinal chi square statistics and FDR-corrected p-values, black points represent distinct non-targeting sgRNAs. (D) Scatterplot of proportion of faint versus punctate VP35 staining patterns of cells for each gene, black points denote individual non-targeting sgRNAs. (E) Significantly enriched gene ontology terms for sets of genes with a high proportion of faint cells (purple) or punctate cells (green). (F) Images of VP35 protein expression (red) overlaid with nuclear mask (gray) in randomly selected cells with sgRNAs targeting the indicated genes. (G) Fine-tuned autoencoder embedding visualized using PHATE for cells with indicated gene knockouts; colors indicate the class labels predicted by the neural network. (H) Ebola virus infection and cell survival (normalized nuclei count relative to untreated) after 48 hours in HeLa cells treated with terpestacin.

While the most significant genes were the same as those identified by the simpler intensity metric (e.g. the intracellular receptor NPC1, the HOPS complex, and negative regulators MESDC2 and ITGB1), the fine-tuned autoencoder model allowed separation of cells that would otherwise have similar protein staining intensities but instead had distinct diffuse, faint, or punctate VP35 protein expression patterns (**Fig. 2d**). Genes with a high proportion of faint cells included many known entry or early infection regulators such as NPC1, as well as genes not previously identified as regulators of infection such as PIK3C3 and members of the GET complex that we had identified using VP35 intensity alone (**Fig. 2d-e**, protein insertion into ER membrane, purple). In contrast, many of the genes associated with a significant increase in punctate VP35 localization have not been previously linked to EBOV infection or to regulation of post-entry viral replication (**Fig. 2e**, green). Key enzymes in *de novo* pyrimidine biosynthesis (DHODH, CAD, and UMPS) previously shown to positively regulate EBOV replication (Luthra et al., 2018; Martin et al., 2018) were significantly enriched for the punctate phenotype, indicating that upon knockout of these genes EBOV had infected the cell and begun formation of inclusion bodies but progression to the next replication step was impeded. Indeed, CAD was previously shown to colocalize with EBOV NP in inclusion bodies, affecting genome replication and transcription (Brandt et al., 2020). In addition to pyrimidine biosynthesis genes, we also identified genes involved in purine biosynthesis (MTHFD1, PAICS, PPAT, ATIC, and GART) not previously associated with EBOV replication, although mycophenolic acid, a small molecule inhibitor of GTP biosynthesis, is known to inhibit EBOV infection (Edwards et al., 2015). MTHFD1 knockdown and treatment with the MTHFD1 inhibitor carolacton was reported to inhibit Zika virus (ZIKV), mumps virus (MuV), and SARS-CoV-2 replication without affecting cell entry, suggesting a common requirement for RNA virus replication and the ability of our deep learning approach to separate effects of genetic knockouts by viral replication stage (Anderson et al., 2021). Knockdown of Elongator complex genes (ELP2-6) also led to an increase in punctate VP35 (**Fig. 2e**). The Elongator complex modifies tRNA molecules needed for translational efficiency and may also be required for optimal viral replication past the inclusion body formation stage (Hawer et al., 2018).

Another major category of genes that elevated VP35 puncta when knocked out were mitochondria-related genes including mitochondrial ribosomes, mitochondrial tRNA synthetases, and mitochondrial respiratory chain complex III and IV members. Specifically, knockouts of UQCRB, a complex III subunit, and HARS2, a mitochondrial tRNA synthetase (**Fig. 2f, g**), resulted in the most enhanced punctate phenotype. Mitochondrial function has not been previously linked to EBOV infection; however, EBOV VP30 and VP35 proteins physically interact with the mitochondrial ribosome and inner membrane components (Batra et al., 2018), and oxidative phosphorylation and expression of mitochondrial translation genes was found to increase upon EBOV infection (Woolsey et al., 2019). Many of the mitochondrial genes identified in our screen have a role in oxidative phosphorylation (OXPHOS), either directly (such as the complex III and IV subunits) or less directly (Arroyo et al., 2016) and OXPHOS plays a role in the propagation of other RNA viruses such as IAV (Bercovich-Kinori et al., 2016). In order to further investigate the effect of UQCRB inhibition on EBOV infection, we treated EBOV-infected HeLa cells with terpestacin, a small molecule inhibitor of UQCRB (Jung et al., 2010), and observed clear reductions in EBOV infection (**Fig. 2h**) with little effect on cell viability as measured by nuclear count at the infection IC50 of 5 μM, highlighting an immediate practical application of results from our deep learning approach.

### Phenotypic profile clustering and matching reveal relationships between modulators of Ebola virus infection

We next more closely examined the 998 genes that significantly altered overall virus infection as measured by VP35 protein or RNA levels (**Table S1**). For each autoencoder model, we extracted embeddings for individual cells and calculated cumulative delta AUCs for each embedding feature between cells with a given sgRNA and non-targeting control cells (**Tables S5**, **S6**). We then clustered the resulting embeddings to identify genes with similar effects (**Fig. 3a**). While the clusters resulting from the two autoencoder models are similar (**Fig. 3b**), the fully unsupervised autoencoder model resulted in a larger number of clusters which were significantly enriched for at least one ontology term using Enrichr (23 versus 7). Single-cell images for genetic knockouts from distinct clusters are shown in **Fig. 3c**. Some genes that scored significantly for increased VP35 protein expression, such as integrins ITGAV (**Fig. 3c**) and ITGB1 (**Fig. S3b**), showed increased intensity in channels not directly related to VP35, such as vimentin and LAMP1, as well as cell rounding. We anticipate that the change in VP35 expression pattern for such genes represents impact on cell morphology or cell health resulting in redistribution of the marker, rather than a change primarily affecting virus replication.

**Figure 3.**
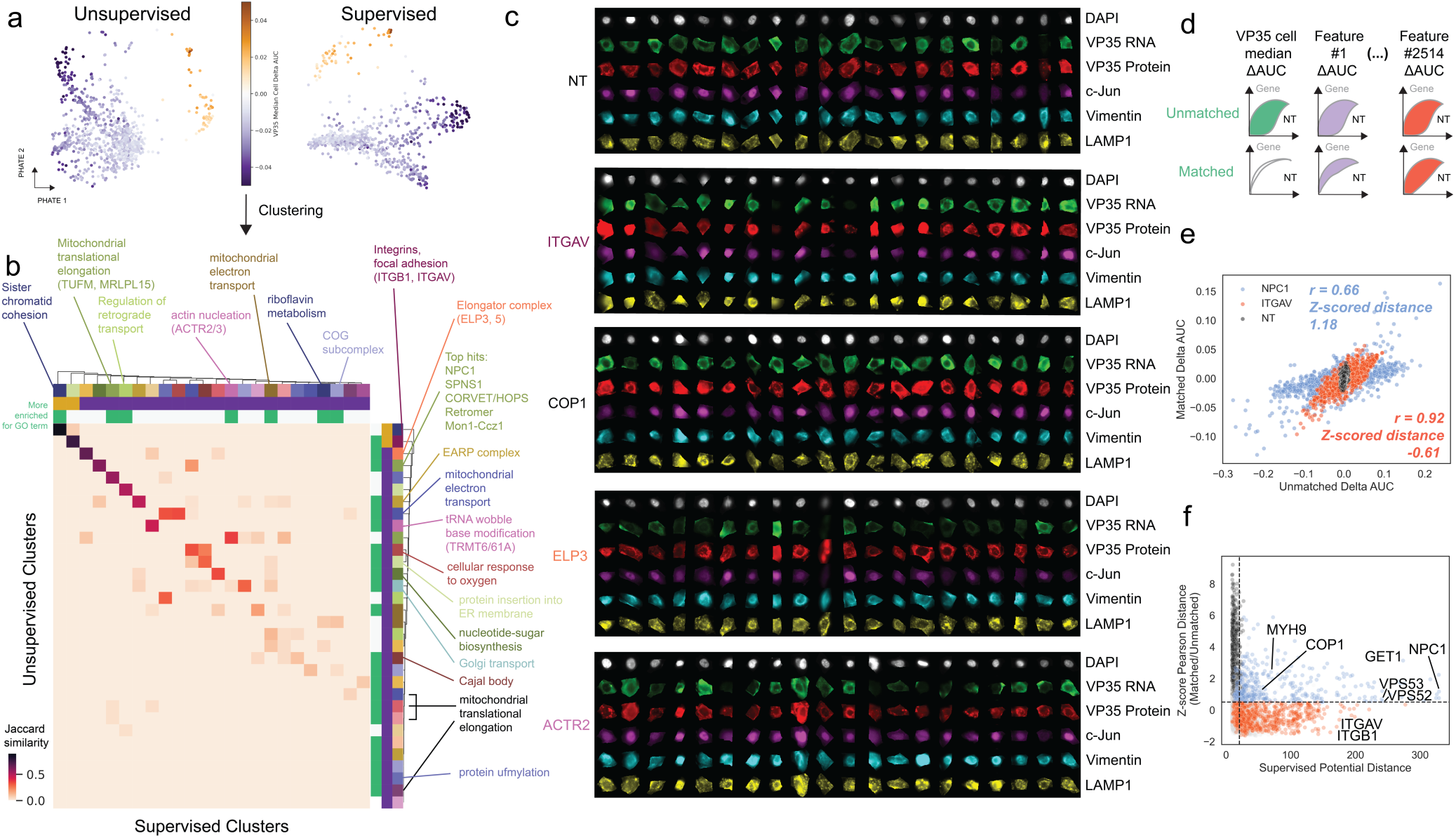
Phenotypic profile clustering and matching on infection level reveal relationships between Ebola virus infection modulators. (A) PHATE visualization of the embeddings derived from the fully unsupervised and the fine-tuned autoencoder models; each point represents the average embedding of a gene knockout of interest. Points in purple indicate genes that decreased Ebola protein intensity upon knockout, while points in gold increased intensity. (B) Heatmap of Jaccard similarities between cluster memberships for all clusters with at least one significant GO term identified using the fully unsupervised model as well as the fine-tuned model. (C) Single-cell images of representative gene knockouts from distinct clusters. (D) Schematic of recalculation of cumulative delta AUCs for autoencoder embeddings between each sgRNA and non-targeting control cells after matching on VP35 protein levels in non-targeting controls. (E) Correlations between delta AUCs calculated without matching on VP35 protein levels compared to with matching on VP35 protein levels for non-targeting controls, NPC1 knockout, and ITGAV knockout cells; each point represents one individual feature. (F) Correlation between the mean PHATE potential distance from non-targeting controls in the fine-tuned autoencoder embedding (indicative of the EBOV infection phenotype strength) and the z-scored Pearson correlation distance between matched and unmatched features from (E); each point represents a gene of interest. Points in red indicate genes that have abnormal cell health, while points in blue represent hit phenotypes under conditions where overall cell health was less affected.

To prioritize genes for further follow-up experimentation, we sought to control observed effects for the level of viral protein expression as a proxy for the level of infection in each cell. For each sgRNA in our screen we sampled non-targeting control cells whose distribution of VP35 protein median intensities matched the VP35 protein median intensity for the sgRNA of interest (**Fig. 3d**), and re-computed delta AUCs between each sgRNA and the matched non-targeting control cells (**Table S7**). Next, we plotted the correlation of delta AUCs for features from all six imaging channels computed without matching (**Table S1**) against those from the matched condition (**Fig. 3e**, **Table S7**). If the primary phenotypic effect of the genetic perturbation is to alter viral infection levels, we expect that the matched AUCs will be close to 0, as the perturbed cells would resemble non-targeting cells with the same infection levels without further morphological changes. Therefore, Pearson correlation between matched and unmatched delta AUCs will be modest, since single-cell images of the genetic perturbation of interest closely match images of non-targeting control cells with a similar infection level. Indeed, we see that features for NPC1 knockout cells are much more modestly correlated between matched and unmatched conditions than features of the integrin ITGAV, which affects global cell morphology rather than viral replication specifically (**Fig. 3e**). Using this approach, we calculated Pearson correlations between matched and unmatched conditions for each of the genes in our screen (**Fig. 3f**) and used this metric to select genes with primary effects on viral replication rather than on cell health for targeted follow-up screening.

### Targeted image-based genetic screens identify cell-and virus-specific filovirus regulators

To evaluate the contributions of cell type, filovirus strain and infection timing to outcomes of the screen we targeted 113 hit genes (manually sub-selected from hits with VP35 protein intensity FDR-adjusted p-value <1e-4, as well as genes with lower impact on cell health, **Fig. 3f**) of interest from our genome-wide screen with 6 sgRNAs per gene and 475,000 to over 1 million cells per condition. Screen conditions varied: 1) timepoints (16 and 24 hours), both earlier than the primary screen (48 hours) to increase power for identification of negative regulators of viral replication; 2) cell lines, the HeLa cells used in the genome-wide screen and Huh7 cells, a liver cell line relevant to filovirus disease; and 3) distinct filoviruses from the genera *Orthoebolavirus* (Ebola virus Mayinga, Sudan virus Gulu) and *Orthomarburgvirus* (Marburg virus Musoke) (**Fig. 4a**, **Table S8**). As in our primary screen, we assayed filovirus protein (VP35 for EBOV and SUDV; VP40 for MARV), and VP35 RNA via probes designed against the VP35 sequences of each virus, as well as c-Jun nuclear translocation.

**Figure 4.**
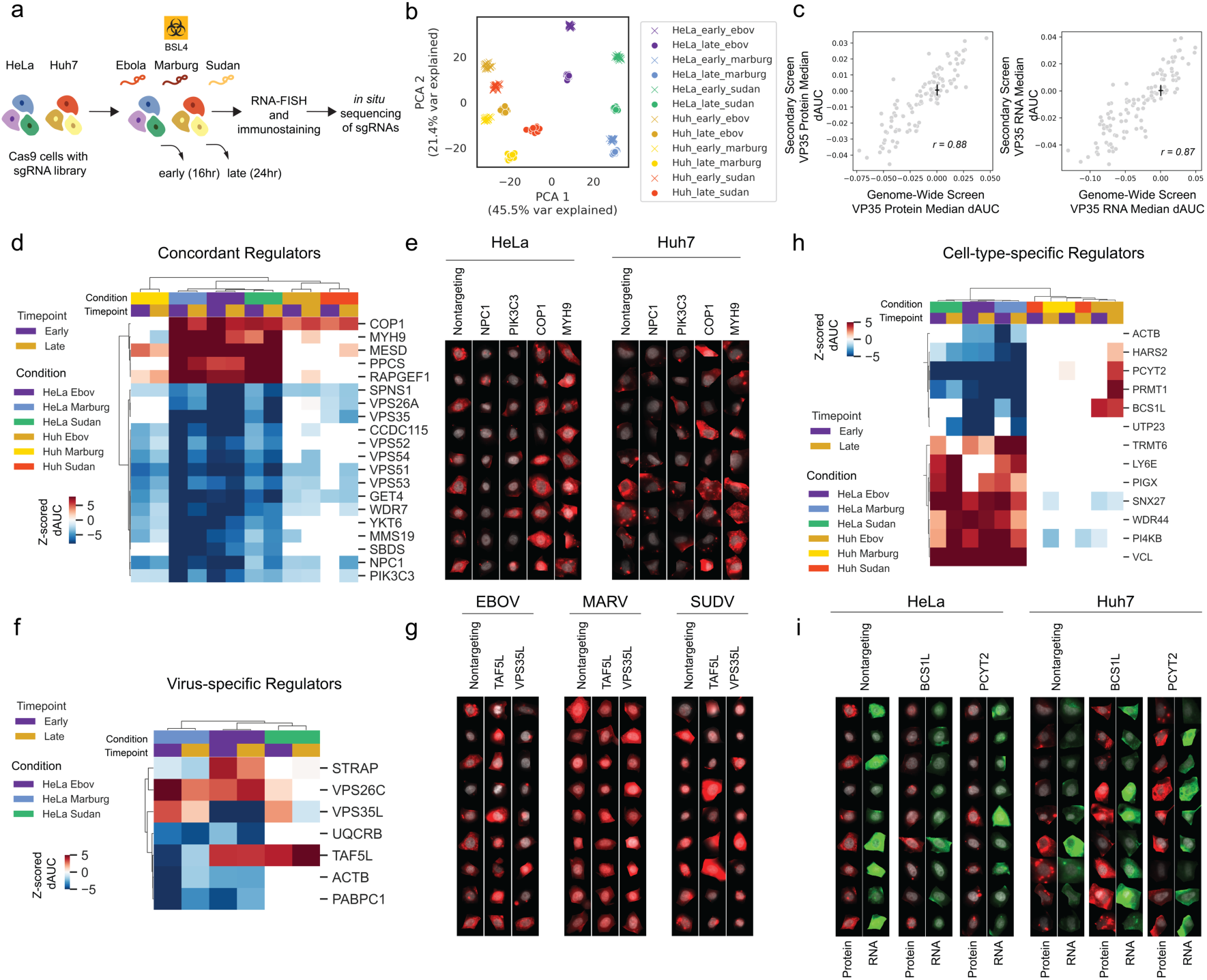
Targeted follow-up screens identify concordant and cell-and virus-specific ebolavirus regulators. (A) Workflow for targeted secondary screens with 2 cell lines, 3 viruses, and 2 timepoints for a total of 12 screening conditions. Phylogenetic relationship modified from previous work (Carroll et al., 2013), scale is substitutions per site. (B) PCA of non-targeting control sgRNA phenotypic profiles from each of indicated screening conditions. Early and late indicate 16 and 24 h infection times respectively. (C) Correlation between genome-wide and secondary screen VP35 protein and RNA delta AUCs (dAUCs); black lines indicate standard deviation for non-targeting control sgRNAs in each screen centered around the mean value for non-targeting sgRNAs in the screen. (D) Heatmap showing z-scored dAUC values for genes concordant across screen conditions (white cells indicate conditions where p > 0.05 relative to non-targeting controls in the same condition). Z-scores were calculated on delta AUC values for all genes in each screen condition relative to means and standard deviations for non-targeting sgRNAs. Hierarchical clustering performed using Pearson correlations. (E) Single-cell images from the secondary screen of select concordant genes (DAPI in gray, VP35 protein in red). (F) Heatmap as in (D) for genes with virus-specific effects (white cells indicate conditions where p > 0.05). (G) Single-cell images from the secondary screen of select virus-specific genes (DAPI in gray, VP35 protein for EBOV/SUDV or VP40 protein for MARV in red). (H) Heatmap as in (D) for genes with cell type-specific effects (white cells indicate conditions where p > 0.05). (I) Single-cell images from the secondary screen of select cell type-specific genes (DAPI in gray, VP35 protein in red, VP35 RNA in green).

Principal component analysis (PCA) of features from non-targeting control sgRNAs from each experimental condition showed separation of outcomes by cell type across PC1 (46% variance explained), with time not being a major contributor to outcome variance for the same cell and virus types. Comparing the cumulative delta AUCs from the genome-wide screen with the 24 hour infection timepoint from the secondary screen resulted in strong correlations (Pearson r >= 0.87) for both VP35 protein and RNA levels (**Fig. 4c**), broadly validating these primary screen hits. Known regulators such as NPC1 and SPNS1 were concordant across cell lines, viruses, and timepoints (**Fig. 4d, e**), as well as retromer complex members (VPS26A and VPS35), which were more recently associated with EBOV cell entry (Poston et al., 2022), and associated genes newly identified in this study, including PIK3C3, GET4, and the GARP complex (VPS51, 52, 53, and 54) (**Fig. 4d**). The ubiquitin ligase COP1 scored as a negative regulator in 10/12 screening conditions (**Fig. 4d, e**). The mechanism for COP1’s regulation of EBOV infection is unclear; however, COP1 targets c-Jun for degradation, among other substrates (Migliorini et al., 2011), so increased c-Jun protein levels in COP1 knockout cells may favor filovirus replication.

We next examined virus-specific regulators (**Fig. 4f, g**), identifying knockout of TAF5L as increasing EBOV and SUDV infection levels while decreasing infection levels of the more distantly related MARV (**Fig. 4f, g**). We also found that VPS35L strongly decreased EBOV infection levels but had modest effects on SUDV and MARV infection, while VPS35 knockout reduced replication of all filoviruses and in both cell types (**Fig. 4d**). VPS35L and VPS35 are members of the retriever and retromer endosomal recycling complexes, respectively (McNally et al., 2017). The distinct effects of VPS35 and VPS35L knockouts may indicate different dependencies of filovirus infection on ER recycling and ER-Golgi trafficking through the retromer and retriever complexes.

Finally, we investigated cell-type-specific regulators (**Fig. 4h, i**). PCYT2, a phospholipid synthesis enzyme which positively regulates formation of lipid droplets (Roberts et al., 2022) increased EBOV infection in Huh7 cells, while decreasing infection in HeLa cells. Interestingly, this gene is more highly expressed in Huh7 cells (Barretina et al., 2012), which have higher lipid droplet content than HeLa cells (Monson et al., 2018), indicating that the differences we observed in viral infection could be due to distinct underlying cellular lipid environments. Several other cell-type-specific regulators were related to mitochondrial function (BCS1L, PRMT1, and HARS2, among others) (**Fig. 4h**, **Fig. S4f**). The strong defect in viral replication observed in HeLa cells upon loss of components related to mitochondrial respiration was typically not observed in Huh7 cells.

### Mechanistic characterization of STRAP as a positive regulator of viral replication

The results of the analyses from the primary screen demonstrate that optical pooled screening is a powerful tool to identify host factors regulating EBOV infection. Secondary screening validated that the hit genes produced robust image-based phenotypes that in many cases were seen across different cell types and filoviruses. One hit that stood out across multiple datasets was Serine/Threonine Kinase Receptor Associated Protein, or STRAP. STRAP was one of only two proteins identified in the random forest analysis as having an unusual phenotype of increased viral RNA relative to VP35 staining intensity upon knockout (**Fig. 1i**) and was identified by the fine-tuned autoencoder model as having a significantly different distribution of VP35 staining phenotypes compared to non-targeting controls (**Fig. 5a**). While STRAP KO did score for decreasing VP35 intensity, it was not among the top hits based on this single measurement. STRAP KO cells showed more than 2-fold increase in the proportion of Faint cells, and a 60% and 50% increase in the proportion of Punctate and Peripheral cells relative to non-targeting controls with an accompanying drop in the proportion of Cytoplasmic cells. Taken together, this profile suggested a unique mechanism of action, where the KO cell population is being shifted into both an earlier stage of infection, represented by the increase in Faint and Punctate classes, and a later stage of infection, represented by the Peripheral class. Examination of single-cell intensity histograms from the genome-wide screen revealed that while there was minimal change in the intensity distribution for protein staining, the RNA levels in STRAP KO cells took on a bimodal distribution, with distinct populations of cells having less viral RNA or more viral RNA compared to the NT controls - an observation only possible at the single-cell level and where single cell-matched perturbation identities were available given the pooled format (**Fig. 5b**). Also of note was a strong reduction in c-Jun nuclear intensity in STRAP KO cells, consistent with previous reports where STRAP plays a role in c-Jun signaling (Reiner et al., 2011).

**Figure 5.**
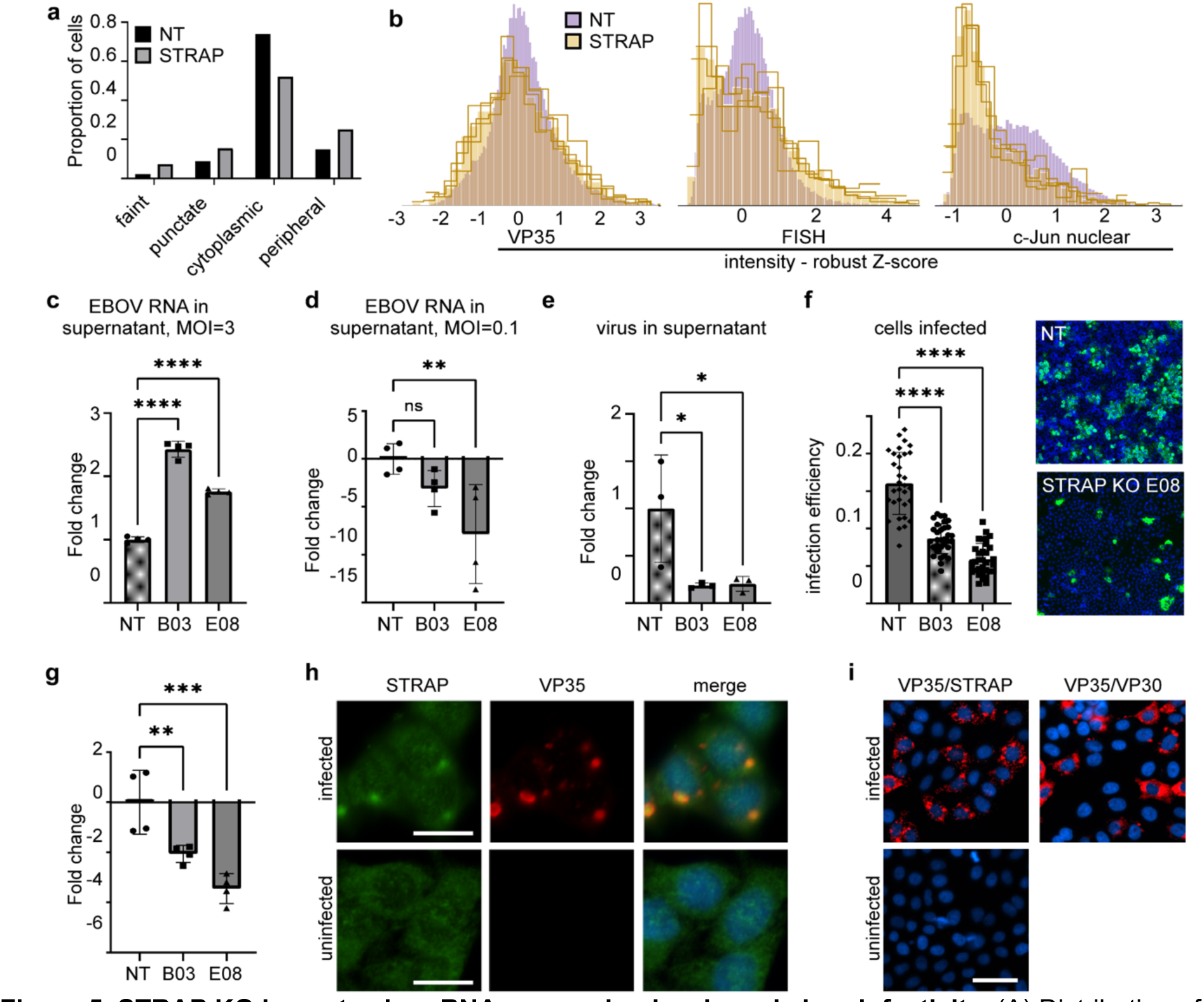
STRAP KO impacts virus RNA expression levels and virus infectivity. (A) Distribution of phenotypes from the primary genome-wide screen as predicted by the fine-tuned autoencoder model for non-targeting controls and STRAP. (B) Histograms of staining intensity for VP35 protein (ICF), VP35 mRNA by FISH, and nuclear c-Jun at single-cell resolution from the primary screen. Each yellow line represents a distinct sgRNA targeting STRAP. (C) Measurement of total viral RNA in clonal cell lines that were KO for STRAP (B03 or E08) or non-targeting cell line (NT) after challenge with virus at MOI = 3 at 72 hours by qPCR. One-way ANOVA comparing each clonal line to the non-targeting control, ****p<0.0001. (D) Measurement of total viral RNA in the supernatant after challenge with MOI = 0.1 after 72 hours by qPCR. **p=0.0086. (E) Measurement of infectious virus in the supernatant after 72 hours by FFU assay, *p<0.025. (F) Measurement of number of cells infected after challenge and allowing 72 h to spread. Infection efficiency was calculated as the number of infected cells divided by total nuclei normalized to the non-targeting control, ****p<0.0001. Right panels show representative images of non-targeting and STRAP knockout cells. (G) Measurements of EBOV negative sense genomic RNA synthesis in cells after 72 hours at MOI =3. One-way ANOVA was used to compare two different STRAP knockout cell lines to non-targeting controls. ***p=0.0002. **p=0.0067. (H) Immunostaining for native STRAP protein and EBOV VP35 in infected cells showing accumulation of STRAP within viral inclusion bodies as marked by VP35 staining. Scale bar is 20 μm. (I) Duolink Proximity Ligation Assay using antibodies against VP35 and STRAP in infected and uninfected cells indicating close association of STRAP with viral proteins within and outside the inclusion bodies. VP35 and VP30 specific antibodies were used on infected cells as a positive control. Scale bar is 50 μm.

To further study the effects of STRAP KO on viral infection, we generated single-cell clones that showed undetectable levels of STRAP by immunoblot, and two were selected for further assays. First, to confirm the phenotype of overall increased RNA in the KOs, cells were incubated with virus at a high multiplicity of infection (MOI = 3) for 72 hours. qPCR of cell lysates showed a strong increase in viral RNA, on average twice that seen in a non-targeting control cell line (**Fig. 5c**). Surprisingly, despite the overall increase in viral RNA in cells, vRNA in the supernatant was reduced by 3-8 fold (**Fig. 5d**) and a 5-fold reduction in infectious virus was observed (**Fig. 5e**). Images of the infected cell monolayer stained for EBOV protein showed a greater number of single cells and smaller foci, indicating that while infection could initiate normally, there was limited virus spread, consistent with cells producing fewer overall infectious virus particles (**Fig. 5f**).

While STRAP KO cells produced fewer infectious virions, the mechanism by which STRAP was modulating viral infection remained unclear. Our previous qPCR assays did not distinguish between positive and negative sense RNA, or between viral mRNA and genomic RNA (gRNA). We used a two-step PCR with primers designed to specifically target the negative sense EBOV gRNA to investigate further. After challenge at an MOI of 3, EBOV gRNA was significantly reduced by 2-3.3 fold in STRAP KO cells compared to the NT controls (**Fig. 5g**). Contrasting with the overall 2-fold increase seen in total viral RNA, the decrease in genomic RNA suggested dysregulation of viral RNA transcription such that the balance between mRNA and genome production had been disrupted.

STRAP is commonly found throughout the cytoplasm and nucleus of the cell, with a diffuse immunofluorescent staining pattern. However, upon EBOV infection, STRAP was recruited to the viral inclusion bodies (**Fig. 5h**) as indicated by STRAP staining directly overlapping VP35, a major component of inclusion bodies. To confirm the close association of STRAP and VP35, we performed a proximity ligation assay (PLA) for STRAP and VP35 (**Fig. 5i**). The PLA assay yields a fluorescent signal when two proteins are within 40 nm of each other (Söderberg et al., 2006), and the intensity and volume of the signal in the cell increases proportionally with a greater number of associative events. Performing PLA with two viral proteins, VP35 and VP30, which would normally closely associate during genome replication, yielded strong signal and displayed both punctate and diffuse phenotypes. PLA performed on uninfected cells showed no signal. When performed with VP35 and STRAP, the signal was similar to that of VP35 and VP30, indicating a similar close association. Punctate and diffuse phenotypes were also visible, suggesting that the interaction can occur outside of virus inclusion bodies. The identification of STRAP as a regulator of the equilibrium between virus RNA and virus protein and how this alters the infectivity of progeny virus, is a new mechanistic dependence that will be further evaluated in future work.

## Discussion

Ebola virus disease remains poorly understood, despite frequent outbreaks with persistent high case fatality rates. Genome-wide screens for host factors impacting EBOV infection have been limited to cytotoxicity-based approaches where cells susceptible to infection succumb to cytopathic effects of the virus. This work has been productive but tends to bias toward strong early blocks to infection, such as the loss of entry receptors. Another approach used more recently for other viruses, such as SARS-CoV-2, has been FACS selection of cells with reporters of infection such as GFP or endogenous virus proteins. The latter approach can identify factors that support infection as well as act as restriction factors. Each approach typically examines one pre-defined parameter of the infection process. Here, we used an image-based pooled screening technology, OPS, to directly link single-cell genetic knockouts with multiple single-cell phenotypes including EBOV infection levels by measuring several distinct aspects of the replication cycle and its effects including viral RNA transcription, protein translation using VP35 expression, and the host responses measured via c-Jun nuclear translocation. The RNA measurements were made possible through a new workflow for integration of targeted RNA FISH with our *in situ* sequencing and immunofluorescence protocols, which enables future image-based screens to integrate host or pathogen RNA levels and localization into cellular phenotypic profiles. By interrogating single-cell images via multiple approaches we have obtained a unique, holistic view of host factor involvement in Ebola virus replication.

The data-rich images obtained report many aspects of the infection process, creating a powerful opportunity to extend our understanding of host factor dependence. We evaluated the data with three complementary analytical approaches and expect the data to be a resource for further future analyses. A first basic analysis separately examined the intensity of each marker of infection (viral protein, RNA, and nuclear c-Jun levels), similar to the analysis procedure for single-phenotype enrichment screens such as FACS screens, with the distinction that each single-cell phenotype is directly linked to the perturbation delivered to that particular cell. This approach revealed many host factors that are known to be important for EBOV infection including NPC1 and members of the HOPS complex, which are all involved in uptake of virus into cells. Unlike previous cell survival screens, we also identified several negative regulators of filovirus replication under multiple conditions, such as COP1, MYH9, and RAPGEF1. Even at this simplistic level, new host factors not previously reported were identified, including PIK3C3, the GARP complex, and the GET complex. We also rigorously validated the effects of host factors on multiple filoviruses and in distinct cell types through a set of 12 secondary screens, which were highly correlated with the primary genome-wide screen (Pearson correlation >= 0.87).

To effectively leverage this high-content imaging data, we next applied deep learning approaches. In particular, we trained unsupervised autoencoders directly on single-cell images, and fine-tuned the resulting model to classify four hand-labeled phenotypes describing distinct categories of VP35 protein subcellular localization. This enabled linkage of altered VP35 punctate staining patterns to specific host factors that likely regulate genome replication, such as some members of the pyrimidine and purine biosynthesis pathways, many of which were not previously linked to Ebola infection and are druggable. Furthermore, terpestacin, a small molecule inhibitor of UQCRB, a gene that our fine-tuned autoencoder model predicted as increasing the fraction of punctate EBOV VP35, is able to decrease EBOV infection in cells, presenting an actionable new therapeutic hypothesis for EBOV infection therapy.

The third key analytical approach we employed was random forest regression analysis to identify host factors that, when knocked out, altered relationships between readouts of infection that are typically correlated. Using this approach, we revealed that knockout of STRAP, a spliceosome component, did not alter levels of virus protein production but rather resulted in increased levels of viral positive-sense RNA and a loss in negative stranded genomes. Furthermore, STRAP was found localized within 40 nm of EBOV VP35, which is involved in the switch between mRNA and gRNA synthesis, in what appeared to be inclusion bodies, indicating that is likely sequestered with the viral replication machinery where it can impact when negative strand synthesis occurs. Since particles of EBOV can assemble in the absence of genomic RNA, adopting other cellular RNAs, (Warfield et al., 2003) our result indicates that virus particle production would be normal but since they lack genomic RNA, they are non-infectious. This outcome represents a new potential target for therapeutic intervention that will be evaluated in future work. Identifying this factor and its mechanism of action would have been challenging through previously established FACS or cytotoxic screening approaches.

High-content single-cell screens with multi-dimensional outputs are becoming routine and were demonstrated at genomic-scale perturbations with a variety of imaging and even transcriptomic readouts (Carlson et al., 2023; Ramezani et al., 2023; Replogle et al., 2022). However, aside from this study and a previous OPS of Sendai virus infection (Carlson et al., 2023), high-content profiling screens of host-pathogen interactions have been limited. Notable recent examples include targeted transcriptome screens that revealed perturbation-induced changes in transcriptional trajectories upon HCMV (Hein & Weissman, 2022) and SARS-CoV-2 (Sunshine et al., 2022) infection. Importantly, high-content perturbation screens like these and that which we report here did not require *a priori* specification of phenotypes of interest but rather resulted in rich data resources that can be mined to rapidly generate and investigate hypotheses based on primary screening data alone (Bock et al., 2022), provided the achievement of adequate quantitative performance of the perturbation and phenotyping steps, and adequate cell sampling. Here, we sampled an average of 2000 cells per gene and key cell feature scores were correlated across independent replicates at r = 0.7-0.9.

Our image-based genetic screens and analytical approaches revealed multiple genes that modulate distinct stages of the EBOV life cycle, from viral entry, to EBOV RNA replication and the formation of viral inclusion bodies, including several genes with available small molecule ligands of potential therapeutic value. This work also serves as a general framework for systematically identifying regulators of distinct steps in host-virus interaction dynamics directly from data produced in a single genetic screen, enabling rapid identification of a broader set of potential therapeutic targets for diverse pathogens.

## Methods

### Library cloning, lentivirus production, and transduction

Libraries were cloned into a CROP-seq-puro-v2 (Addgene #127458) backbone and lentivirus was then produced and transduced as previously described (Feldman et al., 2022). Multiplicity of infection for library transductions was estimated by counting colonies following sparse plating and antibiotic selection with puromycin.

### Virus infection, phenotyping, and *in situ* sequencing for genome-wide screen

HeLa-TetR-Cas9 clonal cells previously described (Feldman et al., 2019) were used for primary screening. Following transduction, cells were selected with puromycin (1 μg/mL) for 3 days after transduction and library representation was validated by NGS. Cas9 expression was induced with 1 μg/mL doxycycline for 1 week and cells were then seeded in ten 6-well glass-bottom dishes at 400,000 cells/well two days prior to fixation. Zaire ebolavirus (strain Mayinga) was added at a multiplicity of infection (MOI) of ∼3 in 2mL media/well for 28 hours prior to fixation.

Cells were fixed by removing media and adding 10% neutral-buffered formalin (Fisher Scientific LC146705) for >6 hours. Cells were permeabilized with 100% methanol for 20 minutes; subsequently, the permeabilization solution was exchanged with PBS-T wash buffer (PBS + 0.05% Tween-20) by performing six 50% volume exchanges followed by three quick washes. Cells were incubated for 30 minutes at 37°C with homemade probe hybridization buffer (30% formamide 5x SSC 0.1% Tween), then incubated with primary probes against VP35 positive-sense RNA (purchased from Molecular Instruments and diluted 1:250 in probe hybridization buffer with 1:100 Ribolock) at 37°C for 4 hours. Samples were then washed 4x5 minutes in probe hybridization buffer at 37°C, washed 3x in PBS-T, and incubated in reverse transcription mix at 37°C overnight. The reverse transcription mix consisted of 1x RevertAid RT buffer, 250 μM dNTPs, 0.2 mg/mL BSA, 1 μM RT primer with LNA bases, A+CT+CG+GT+GC+CA+CT+TTTTCAA, 0.8 U/μL Ribolock RNase inhibitor, and 4.8 U/μL RevertAid H minus reverse transcriptase in 750 μL/well. After reverse transcription, cells were washed 5x with PBS-T and post-fixed using 3% paraformaldehyde and 0.1% glutaraldehyde in PBS for 30 minutes, followed by washing with PBS-T 3 times. Samples were then incubated in homemade HCR FISH amplification buffer (5x SSC 0.1% Tween) at room temperature for 30 minutes. Meanwhile, HCR hairpins (Molecular Instruments B1 probes conjugated to Alexa Fluor 488) were separately prepared by heating at 95°C for 90 seconds and then cooling to room temperature in the dark for 30 minutes. Next, samples were incubated with probes diluted 1:125 in probe amplification buffer for 2 hours at room temperature. Following incubation, excess hairpins were removed by washing 5 times for 5 minutes with probe amplification buffer. Primary antibodies against VP35 (1:3200 dilution), c-Jun (1:1800 dilution, Cell Signaling Technology Cat# 9165, RRID:AB_2130165), and vimentin (1:1300 dilution, Abcam Cat# ab24525, RRID:AB_778824) were added by incubating samples for 3.5 hours at 37°C in 3% BSA (VWR Cat# 97061-422) in PBS. Samples were then washed 3x in PBS-T for 3 minutes, and incubated with secondary antibodies: 1:1800 donkey anti-mouse antibody (Jackson ImmunoResearch Labs Cat# 715-006-151, RRID:AB_2340762) disulfide-linked to Alexa Fluor 594 (Thermo Fisher A10270) via a custom conjugation, 1:1800 donkey anti-rabbit antibody (Jackson ImmunoResearch Labs Cat# 711-006-152, RRID:AB_2340586) disulfide-linked to Alexa Fluor 647 (Thermo Fisher Scientific A10277) via a custom conjugation, and goat anti-chicken DyLight 755 (Thermo Fisher Scientific SA5-10075), in 3% BSA for 3 hours at 37°C. Finally, samples were washed 6x with PBS-T for 3 minutes each, and 2x SSC with 200ng/mL DAPI was added to visualize nuclei to image samples. Following imaging, Alexa Fluor 594 and 647 antibodies were destained with 50mM TCEP in 2x SSC for 45 minutes at room temperature and FISH signal removed through treatment with 80% formamide in 2x SSC for 30 minutes at room temperature. In situ amplification of sgRNA sequences was then completed by incubating samples in a padlock probe and extension-ligation reaction mixture (1x Ampligase buffer, 0.4 U/μL RNase H, 0.2 mg/mL BSA, 100 nM padlock probe -/5Phos/GTTTCAGAGCTATGCTCTCCTGTTCGCCAAATTCTACCCACCACCCACTCTCCAAAGGACGAAACACCG, 0.02 U/μL TaqIT polymerase, 0.5 U/μL Ampligase and 50 nM dNTPs) for 5 minutes at 37°C and 90 minutes at 45°C, and washing 2 times with PBS-T. Circularized padlocks were amplified using a rolling circle amplification mix (1x Phi29 buffer, 250 μM dNTPs, 0.2 mg/mL BSA, 5% glycerol, and 1 U/μL Phi29 DNA polymerase) at 30°C overnight. AlexaFluor 488-conjugated LAMP1 (Cell Signaling Technology Cat# 58996, RRID:AB_2927691) was visualized via incubation for 2 hours at 37°C in 3% BSA at 1:500 dilution. Following imaging, in situ sequencing was performed as previously described using sequencing primer GCCAAATTCTACCCACCACCCACTCTCCAAAGGACGAAACACCG for 12 cycles (Feldman et al., 2019).

### Virus infection, phenotyping, and *in situ* sequencing for secondary screen

HeLa-TetR-Cas9 clonal cells used in the genome-wide screen and polyclonal Huh7 cells transduced with 311-Cas9 (Plasmid Addgene #96924) and selected using 10 µg/mL blasticidin for 7 days were used for secondary screening. EBOV (strain Mayinga), Marburg virus (strain Musoke), Sudan virus (strain Gulu) were added at MOIs of ∼3 in 2mL media/well for 16 hours or 24 hours prior to fixation. Viral RNA was amplified as described for the primary screen using virus strain-specific probes purchased from Molecular Instruments for positive-sense VP35. Antibody staining and in situ sequencing was performed as described for the primary screen except for Marburg virus, where an antibody against VP40 (1:1000, Integrated BioTherapeutics 0203-012) rather than VP35 was used since no antibody for Marburg VP35 was available. In addition, antibodies for vimentin and LAMP1 were omitted to allow for slower higher-magnification imaging. For Sudan virus, the same VP35 antibody was used as it recognized both EBOV and Sudan VP35 protein. *In situ* sequencing was performed for 6 cycles.

### Fluorescence microscopy for primary genome-wide and secondary targeted follow-up screens

All *in situ* sequencing images were acquired using a Ti-2 Eclipse inverted epifluorescence microscope (Nikon) with automated XYZ stage control and hardware autofocus. The Lumencor CELESTA Light Engine was used for fluorescence illumination and all hardware was controlled using NIS elements software with the JOBS module. *In situ* sequencing cycles were imaged using a 10X 0.45 NA CFI Plan Apo λ objective (Nikon MRD00105) with the following filters (Semrock) and exposure times for each base: G (546 nm laser at 40% power, emission 575/30 nm, dichroic 552 nm, 200 ms); T (546 nm laser at 40% power, emission 615/24 nm, dichroic 565 nm, 200 ms); A (637 nm laser at 40% power, emission 680/42 nm, dichroic 660 nm, 200 ms); C (637 nm laser at 40% power, emission 732/68 nm, dichroic 660 nm, 200 ms). For the genome-wide primary screen, phenotyping images were acquired using a 20X 0.75 NA CFI Plan Apo λ objective (Nikon MRD00205) with the following filters (Semrock unless otherwise noted) and exposure times: DAPI (405 nm laser at 5% power, Chroma Multi LED set #89402, 50ms), AF488 (477 nm laser at 30% power, Chroma Multi LED set #96372, 200ms), AF594 (546 nm laser at 10% power, emission 615/24 nm, dichroic 565 nm, 200ms), AF647 (637 nm laser at 10% power, emission 680/42 nm, dichroic 660 nm, 200ms), Dylight 755 (749 nm laser at 10% power, emission 820/110 nm, dichroic 765 nm, 200ms).

For the secondary screen, phenotyping images were acquired using a 40x 0.95 NA CFI Plan Apo λ objective (Nikon MRD70470) with the following filters and exposure times: DAPI (405 nm laser at 5% power, Chroma Multi LED set #89402, 50ms), AF594 (546 nm laser at 10% power, emission 615/24 nm, dichroic 565 nm, 200ms), and AF647 (637 nm laser at 10% power, emission 680/42 nm, dichroic 660 nm, 200ms).

### Quantification and Statistical Analysis

#### Image analysis

Images of cell phenotype and *in situ* sequencing of perturbations were manually aligned during acquisition using nuclear masks to calibrate the plate position to each of the four corner wells during screening. Alignment was then refined computationally via cross-correlation of DAPI signal between imaging acquisitions. Nuclei and cells were detected and segmented as previously described and *in situ* sequencing read calling was performed as previously described (Feldman et al., 2022). Data analysis functions were written in Python, using Snakemake for workflow control (Köster & Rahmann, 2012). Image analysis code is available on GitHub. Briefly, for segmentation of phenotyping images from the primary screen, nuclei were segmented using the following parameters: nuclei smooth = 4, nuclei radius = 15, nucleus area 90-1200, DAPI intensity threshold = 1350. Cells were segmented using signal in the vimentin channels, at an intensity threshold = 3000. For segmentation of *in situ* sequencing images from the primary and secondary screens, nuclei were segmented using the following parameters: nuclei smooth = 1.15, nuclei radius = 15, nucleus area 20-400, DAPI intensity threshold = 1000-2000, adjusted differently for each plate. Cells were segmented using signal in the four sequencing channels at intensity thresholds adjusted for each plate, between 2500 and 4200. For segmentation of phenotyping images from the secondary screen, nuclei were segmented using the following parameters: nuclei smooth = 9, nuclei radius = 100, nucleus area 200-18,000 for HeLa cells or 200-50,000 for Huh7 cells, DAPI intensity threshold = 4000 for HeLa cells and 3000 for Huh7 cells. Cells were segmented using background cell signal in the Jun channel, at an intensity threshold = 1525-1900 for Hela cells (depending on the assay plate) and 1625-1825 for Huh7 cells. All other parameters used for analysis were set to default settings.

### Optical pooled screen analysis

Only cells with a minimum of one read matching a barcode in the library were analyzed. For the genome-wide screen, only genes with a minimum of one read matching an sgRNA in the library and 2 sgRNAs with at least 50 cells/sgRNA were considered for analysis. Features were normalized on a per-cell basis relative to cells in the same field of view by subtracting the median and dividing by the MAD x 1.4826 (Bray et al., 2016) and scores for features relative to non-targeting controls were determined by calculating differences in cumulative AUCs. These delta AUCs were averaged over sgRNAs for a given gene and significance was determined by comparing delta AUCs for individual sgRNAs to distributions bootstrapped from non-targeting control cells (bootstrapped 100,000 times). Gene-level p-values were calculated using Stouffer’s method and then corrected using the Benjamini-Hochberg procedure. Random forest regression models were trained on features from the VP35 protein channel only (for predicting VP35 RNA FISH levels) or the VP35 protein and RNA levels (for predicting c-Jun nuclear intensity) for 50,000 randomly selected non-targeting control cells using sklearn.ensemble.RandomForestRegressor with scikit-learn 1.1.3, random state set to 7, n_estimators = 100, and max_features = ‘sqrt’ (Pedregosa et al., 2011). Statistical significance was determined as described for the cumulative AUC analysis above.

### Dimensionality Reduction, Clustering, and Gene Enrichment Analysis

In Figures 1 and 2, Enrichr results (Kuleshov et al., 2016) were determined using gseapy 0.14.0 (Z. Fang et al., 2023) with the 2021 KEGG and GO gene sets and 2016 Reactome gene sets. PHATE 1.0.10 (Moon et al., 2019) was used to perform dimensionality reduction on single cells with Euclidean distance, cosine mds distance, gamma = 1, knn = 5, 20 PCs in Figure 2. For gene-level clustering in Figure 3, PHATE with Euclidean distance, cosine mds distance, gamma = 1, knn = 3 and the number of PCs giving 95% of the variance (282 for unsupervised and 372 for supervised features) were used. Following dimensionality reduction, Leiden clustering was performed with the resolution parameter determined based on the Adjusted Rand Score (ARS) computed by subsampling 95% of the data and re-clustering at least ten times for each resolution value.

### Deep learning model

#### Unsupervised Training

First, we trained an unsupervised convolutional autoencoder on 40 million 64x64 six-channel single-cell images. We implemented a U-Net style architecture (Ronneberger et al., 2015) with an encoder containing 5 convolutional layers and a decoder containing 5 convolutional layers (Fig. 2a). We used strided convolution to reduce dimensionality of the images in the encoder, resulting in a 2048-dimensional embedding. Bilinear upsampling was used in the decoder to map from this 2048-dimensional latent space back to the original image space.

For model selection, we used an 80/20 train/test split. In each epoch of training, at least 256 cells were sampled from each training field of view. As a loss function, we used the mean squared error (MSE) over all reconstructed pixels within the watershed mask for each cell of interest. We used the Adam optimizer with a learning rate of 0.001, trained for 50 epochs with a random seed. Training and test loss curves are shown in Figure S2a.

#### Supervised Training

We then fine-tuned the encoder of the trained autoencoder by training a classification head with the Negative Log-Likelihood loss (NLL Loss) using Adam with a learning rate of 0.001. We used 3,889 cell images manually labeled with one of four phenotypic categories in the set {faint, punctate, cytoplasmic, peripheral}. We used stratified dataset splitting implemented in scikit-learn (Pedregosa et al., 2011) to ensure a balanced distribution of phenotypes across classes. Namely, we reserved 25% of the dataset, or 973 cell images, as a held-out test set, and the remaining 2,916 images were split into 4 stratified folds (train/validation) and used for fine-tuning the autoencoder.

For fine-tuning the autoencoder, we applied the following data augmentations using the available image transforms in PyTorch: (1) rotations up to 180 degrees; (2) random vertical and horizontal flips; (3) random perspective with a distortion scale of .1; (4) random affine transformations with a shear of 10 and a scale of (.75, 1.25); and (4) Gaussian blur with a kernel size of 5 and standard deviation uniformly selected in the interval (0.05, 0.5). For training our model, we constructed a batch by sampling 5 images from each of the four classes, applying 5 random transformations to each image, resulting in 100 images per batch. We sampled 100 batches in any given epoch. The balanced sampling strategy was used to account for any class imbalances. We trained for a total of 50 epochs and chose the model that achieved the highest balanced accuracy defined as accuracy across each class weighted by the proportion of validation samples in the class.

To fairly compare model performance across embeddings from various standard deep learning models on the held-out test set, we trained Support Vector Machines (SVMs) with a linear kernel on the embeddings extracted from these models on the held-out test set In particular, we split our test embeddings according to a 95-5 split, trained the SVM on the 95%, and evaluated model performance on the remaining 5%. We trained SVMs on the embeddings obtained from the fully unsupervised autoencoder model, the fine-tuned autoencoder model, a pre-trained ResNet model, and hand-crafted features (Fig. S2d).

To determine sgRNAs that significantly altered the proportion of cells in each phenotypic category, an ordinal chi squared test was performed using R 4.2.2 with the coin package (v1.0.9), and the results were combined at the gene level using Stouffer’s method.

### Identification of Terpestacin as an Antiviral Compound

The evening before the infection, ∼15,000 HeLa cells were seeded into each well of a 96 well plate. The following day, terpestacin (Aurora Fine Chemicals, CA, USA), a known small molecule inhibitor of UQCRB, was dosed onto cells in a 9-point, 2-fold dose curve beginning at 50µM. Cells were infected with EBOV at an MOI of 0.1-0.2 and allowed to infect for 48 hours. The plates were then fixed in 10% neutral-buffered formalin for >6 hours and removed from containment. The cells were washed, permeabilized with 0.1% Triton X-100, blocked with 3.5% BSA, and immunostained with an anti-EBOV GP monoclonal antibody (IBT, MD, USA). After several hours incubating at 37°C, the cells were washed in PBS and incubated in anti-mouse Alexa Fluor 488 secondary antibody (Thermo Fisher, MA, USA). Cells were again washed in PBS and nuclei were counterstained with Hoechst 33342. The plate was imaged with a BioTek Cytation 1 automated plate imager. Images were fed into a custom pipeline in CellProfiler (Broad Institute, MA, USA) used to count the number of infected cells and nuclei. Infection efficiency was calculated as the number of infected cells divided by the total number of nuclei (as a proxy for total cell count) and normalized to the average of cells treated with DMSO only. The total nuclei count, normalized to the average of the negative controls, was used to check for potential cytotoxicity. The normalized infection efficiency and normalized nuclei counts were plotted in GraphPad Prism 8.0.0 (GraphPad Software, CA, USA). A four parameter variable slope nonlinear regression was used to fit the data and calculate the antiviral IC50 of the compound.

### Generation of STRAP KO Cells

The evening before transduction, 250,000 HeLa-TetR-Cas9 cells were seeded into a 12 well plate in DMEM supplemented with 10% FBS. The following morning, the medium was replaced with DMEM containing 8µg/mL polybrene. A lentiviral vector containing an sgRNA targeting STRAP purchased from the Broad Institute’s Genetic Perturbation Platform was added and the cells were spinoculated at 1000xg for 2 hours at 33°C. After spinoculation, the cells were transferred to the incubator for 3 hours, follow by medium replacement. 24 hours after spinoculation, the cells were split into medium containing 2µg/mL puromycin to begin selection. 48 hours after beginning selection, the medium was replaced with that containing 2µg/mL puromycin and 1µg/mL doxycycline to induce Cas9 activation. Once the cells reached confluency, they were split into medium containing only 1µg/mL doxycycline and were maintained in doxycycline for one week. After one week, the cells were split into basic medium and single cell clones were obtained. Clonal populations were identified and verified as being STRAP KOs by capillary electrophoresis using a Jess automated immunodetection machine (Bio-Techne, MN, USA). Two clones, identified as B03 and E08, had no detectable STRAP expression and were used for subsequent experiments.

### qPCR of EBOV RNA

To detect and quantify total cellular EBOV RNA, NT or STRAP KO cells were infected at the indicated MOI and incubated for approximately 16 hours. Cells were washed once with PBS and harvested in TRIzol (ThermoFisher, MA, USA). RNA was processed by a Zymo Direct-zol RNA Miniprep kit as per manufacturer’s instructions. RT-qPCR was performed using the NEB Luna Universal Probe One-Step RT-qPCR kit (E3006L) on a Bio-Rad CFX Opus 96 Real Time PCR system. Primer/probe sequences for EBOV NP (forward-GCAGAGCAAGGACTGATACA, reverse-GTTCGCATCAAACGGAAAAT, probe-FAM-CAACAGCTT-ZEN-GGCAATCAGTAGGACA-IABkFQ) and human GAPDH (forward-ACATCGCTCAGACACCATG, reverse-GTAGTGAGGTCAATGAAGGG, probe-Cy5-AAGGTCGGAGTCAACGGATTTGGTC-IAbRQSp) were previously established. The thermocycler protocol was as follows: 55°C for 10 minutes, 95°C for 1 minute, then 40 cycles of 95°C for 10 seconds and 60°C for 30 seconds. Primers and probes were synthesized by IDT (IA, USA). Four concentrations of synthetic RNA standards were used to calculate genome equivalents from Cq values. Each standard and sample were run in duplicate technical replicates and averaged. To detect extracellular EBOV RNA, NT or STRAP KO cells were infected at the indicated MoIs and incubated for ∼72 hours. Supernatants from infected cells were harvested into TRIzol LS (ThermoFisher, MA, USA) in a 3:1 TRIzol LS to supernatant ratio. RNA was processed and PCR was performed as above using the same primers.

To quantify genome specific EBOV RNA, whole cell lysates of infected NT or STRAP cells were generated and processed as above. A two-step PCR was performed to specifically target EBOV genomic RNA, which has been previously described. For reverse transcription, an Invitrogen SuperScript III Reverse Transcriptase kit was used following the manufacturer’s protocol. Briefly, 4µL purified RNA was mixed with 1µL 2uM gene-specific primer (EBOV-18046 forward, 5’ GAGTTGATTAGTGTGTGCAATAGGTTTAC 3’), 1µL 10mM dNTP mix, and 4µL nuclease-free water. The mixture was denatured for 5 minutes at 65°C, then cooled on ice for greater than 1 minute. After cooling, 4µL First Strand Buffer, 1µL 0.1M DTT, 1µL RNase inhibitor, 1µL SuperScript III reverse transcriptase, and 3µL nuclease-free water were added (total reaction volume of 20µL). Reverse transcription was performed at 55°C for 1 hour, and the enzyme was then inactivated at 85°C for 5 minutes. 4µL of the first reaction was used as the template for the subsequent qPCR using the BioRad iTaq Universal SYBR Green Supermix according to the manufacturer’s protocol.

### EBOV Immunostaining and Infection Efficiency Calculation

Cells were infected as described above with a low MOI. After a 72 hour infection, the plates were fixed in 10% formalin and removed from containment. Cells were immunostained with an anti-EBOV GP mouse antibody (IBT, MD, USA) and Alexa Fluor 488 anti-mouse secondary (Thermo Fisher, MA, USA). Nuclei were counterstained with Hoechst 33342. Images were taken with a Biotek Cytation 1 automated plate imager. The number of infected cells and nuclei were quantified using a customized pipeline on CellProfiler (Broad Institute, MA, USA) and infection efficiency was calculated by dividing the total number of infected cells by the total number of nuclei.

### Focus Forming Unit Assay

Non-targeting and STRAP knockout HeLa cells were infected as described above with a low MOI in triplicate. After a 72 hour infection, supernatant was harvested from samples and diluted out onto HeLa cells in a 96 well plate in a 2-fold series. Cells were incubated for another 72 hours to allow viral replication and spread. The plates were formalin fixed, removed from containment, and immunostained for EBOV GP as above. One focus forming unit was defined as a cluster of >5 infected cells. The lowest dilution with an average of >50 FFUs was used as the endpoint to calculate the viral titers. The data was normalized to the average of the non-targeting controls.

### Co-immunostaining of EBOV VP35 and STRAP Proteins

The night prior to infection, ∼30,000 HeLa cells were seeded into 8 well chamber slides. Cells were infected with EBOV at an MOI of ∼3 for 16-18 hours, then fixed in 10% formalin and removed from containment. Cells were permeabilized in 0.1% Triton X-100, blocked in 3.5% BSA, and immunostained with our anti-EBOV VP35 mouse antibody and an anti-STRAP rabbit antibody (Atlas Antibodies, Bromma, Sweden) at 1:3000 and 1:1000 respectively for several hours at 37°C. Cells were washed and anti-rabbit Alexa Fluor 488 and anti-mouse Alexa Fluor 594 secondary antibodies were added (Thermo Fisher, MA, USA). After 1-2 hours, the cells were washed, and nuclei were counterstained with Hoechst 33342. Images were taken on a Nikon Ti2 Eclipse microscope.

#### Proximity Ligation Assay

The night prior to infection, ∼7,000 HeLa cells were seeded into an 18 well chamber slide (Ibidi, Gräfelfing, Germany). The following evening, cells were infected with EBOV at an MOI of ∼3. After 16-18 hours, the cells were fixed in 10% formalin and removed from containment. Cells were washed thoroughly with PBS, and the Duolink PLA kit was used (Sigma-Aldrich, MO, USA) according to the manufacturer’s protocol. Briefly, the blocking solution was added for 1 hour at 37°C. The anti-EBOV VP35 mouse antibody and anti-STRAP rabbit antibody were diluted 1:3000 and 1:1000 respectively in antibody diluent and incubated at 37°C for 2-3 hours. The cells were washed in wash buffer A and diluted Duolink PLA probes were added for 1 hour at 37°C. Cells were washed in wash buffer A and the ligase was added in ligase buffer for 30 minutes at 37°C. Cells were again washed in wash buffer A and the polymerase diluted in amplification buffer was added for 100 minutes at 37°C. The cells were finally washed in wash buffer B and the nuclei were counterstained with Hoechst 33342. Images were taken on a Nikon Ti2 Eclipse microscope.

## Data and Code Availability

Code is available at https://github.com/beccajcarlson/EBOVOpticalPooledScreen. Imaging data is available on Google Cloud Storage at gs://opspublic-east1/EBOVOpticalPooledScreen.

## Acknowledgments

We thank members of the Blainey, Davey, Uhler, and Hacohen labs for critical feedback and discussions. We thank Celeste Diaz and Julia Bauman in the lab of J.T. Neal at the Broad Institute for assistance in developing custom antibody conjugations. The HeLa cell line was used in this research. Henrietta Lacks, and the HeLa cell line that was established from her tumor cells without her knowledge or consent in 1951, have made significant contributions to scientific progress and advances in human health. We are grateful to Lacks, now deceased, and to the Lacks family for their contributions to biomedical research. This work was supported by the Broad Institute through startup funding (to P.C.B.) and the BN10 program, and two grants from the National Human Genome Research Institute (HG009283 and RM HG006193). P.C.B. was supported by a Career Award at the Scientific Interface from the Burroughs Wellcome Fund. R.A.D. was supported by P01AI120943. R.J.C. was supported by a Fannie and John Hertz Foundation Fellowship and an NSF Graduate Research Fellowship. C.F.B. was supported by R01AI148663 and P01AI120943. A.R. was supported by a George F. Carrier Postdoctoral Fellowship.

## Author Contributions

R.J.C. and J.J.P. designed the approach with input from all authors. R.J.C, J.J.P., B.Y.S., and N.T. performed experiments. R.J.C. performed analysis aside from developing the deep learning model. A.R. and G.S. developed the deep learning methodology and designed the architecture for it with input from R.J.C., J.J.P., and C.U., and G.S. trained the model to obtain the single-cell embeddings. A.S. provided critical feedback and performed custom antibody conjugation. G.A., D.L., C.F.B., N.H., C.U., R.A.D., and P.C.B. supervised the research. R.J.C. and J.J.P. wrote the manuscript with contributions from all authors.

## Competing Interests

P.C.B. is a consultant to or holds equity in 10X Genomics, General Automation Lab Technologies/Isolation Bio, Celsius Therapeutics, Next Gen Diagnostics, Cache DNA, Concerto Biosciences, Stately, Ramona Optics, Bifrost Biosystems, and Amber Bio. His laboratory receives research funding from Calico Life Sciences, Merck, and Genentech for work related to genetic screening. N.H. holds equity in and advises Danger Bio/Related Sciences, owns equity in BioNtech and receives research funding from Bristol Myers Squibb. C.U. serves on the Scientific Advisory Board of Immunai, Relation Therapeutics and Focal Biosciences, and receives research funding from AstraZeneca and Janssen Pharmaceuticals. The Broad Institute and MIT may seek to commercialize aspects of this work, and related applications for intellectual property have been filed. A.S. is an employee at Genentech and R.J.C. is an employee at Flagship Pioneering.

## Supplementary Information

**Table S1.** Per-gene mean cumulative delta AUC scores and p-values for VP35 protein, VP35 RNA FISH, and c-Jun channels in genome-wide screen.

**Table S2.** Per-gene mean cumulative delta AUC scores for all predefined features.

**Table S3.** Per-gene mean random forest regression results for VP35 RNA FISH and c-Jun predictions.

**Table S4.** Per-gene mean deep learning predictions of VP35 subcellular protein localization and associated ordinal chi square statistics and p-values.

**Table S5.** Per-gene mean cumulative delta AUC scores for unsupervised autoencoder features.

**Table S6.** Per-gene mean cumulative delta AUC scores for supervised transfer learned features.

**Table S7.** Mean delta AUCs per gene for all features with or without matching to infection level.

**Table S8.** Per-gene mean delta AUC scores and p-values for VP35 protein and VP35 RNA FISH channels in all secondary screen conditions.

**Figure S1.**
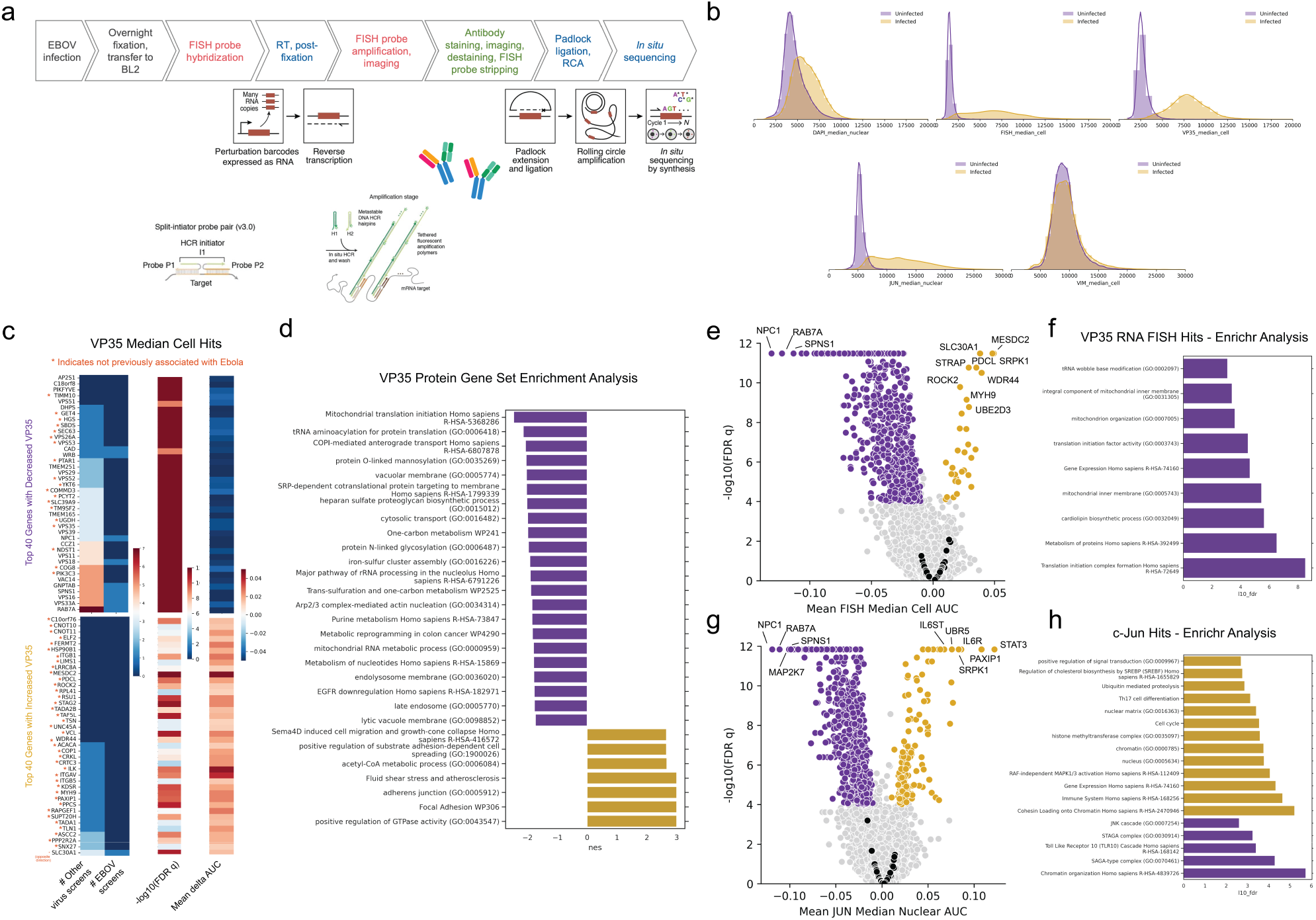
(A) Integration of optical pooled screening workflow with RNA FISH detection using HCR amplification. (B) Histograms of intensity features in five channels for non-targeting controls cells that were infected or not infected in the genome-wide optical pooled screen. (C) Top 40 hits with increased or decreased VP35 protein levels and the number of non-Ebola virus genetic screens or Ebola-specific genetic screens they scored in. Genes not previously associated with Ebola in the literature are marked with an orange asterisk. (D) Gene set enrichment analysis of genes with significantly decreased (purple) or increased (gold) Ebola virus VP35 protein levels. (E) Volcano plot showing genes that scored significantly for changes in VP35 RNA levels by FISH. (F) Enrichr analysis of gene ontology terms significantly enriched in genes that reduced VP35 RNA levels. (G) Volcano plot showing genes that scored significantly for changes in c-Jun levels. (H) Enrichr analysis of gene ontology terms significantly enriched in genes that reduced or enhanced c-Jun levels.

**Figure S2.**
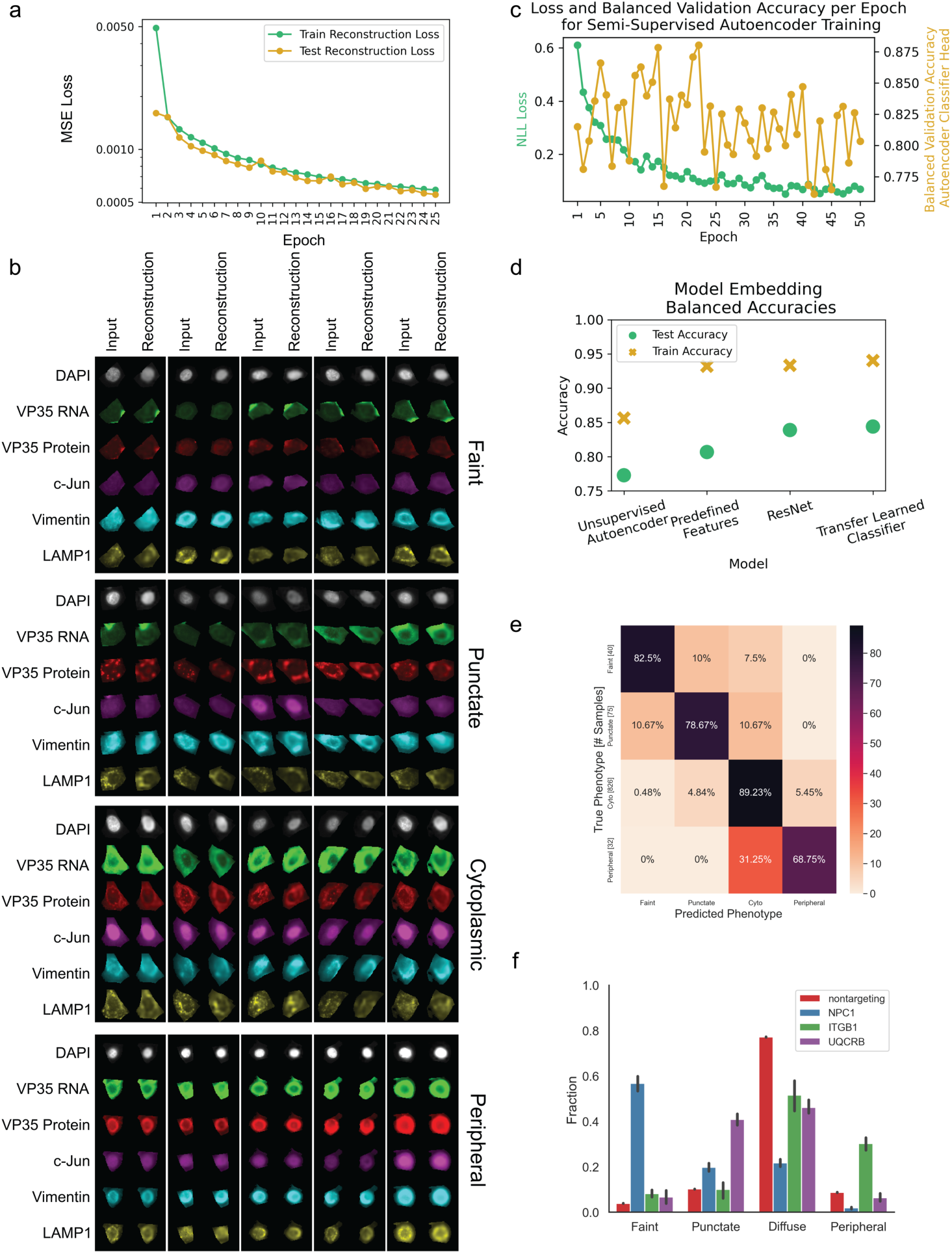
(A) Fully unsupervised autoencoder reconstruction losses for training and test sets across 25 epochs. (B) Examples of manually labeled faint, punctate, cytoplasmic, and peripheral input cell images with accompanying unsupervised autoencoder reconstructions. (C) Fine-tuned autoencoder trained using negative log likelihood loss with balanced validation accuracy also reported across 50 epochs of training. (D) Best model train and test set accuracies for the VP35 protein localization prediction task using SVMs on latent embeddings from the unsupervised autoencoder, predefined features, a Resnet-50 architecture trained on the prediction task, or the fine-tuned autoencoder. Predefined features include intensity, correlation, and texture morphological features similar to those previously described for Cell Painting (Bray et al., 2016). (E) Confusion matrix of model predictions vs manually labeled classifications on model test set. (F) Proportion of cells in each VP35 localization category for non-targeting controls and the genes with the largest proportion of faint (NPC1), punctate (UQCRB), and peripheral (ITGB1) cells. Error bars indicate SEM across sgRNAs targeting the same gene.

**Figure S3.**
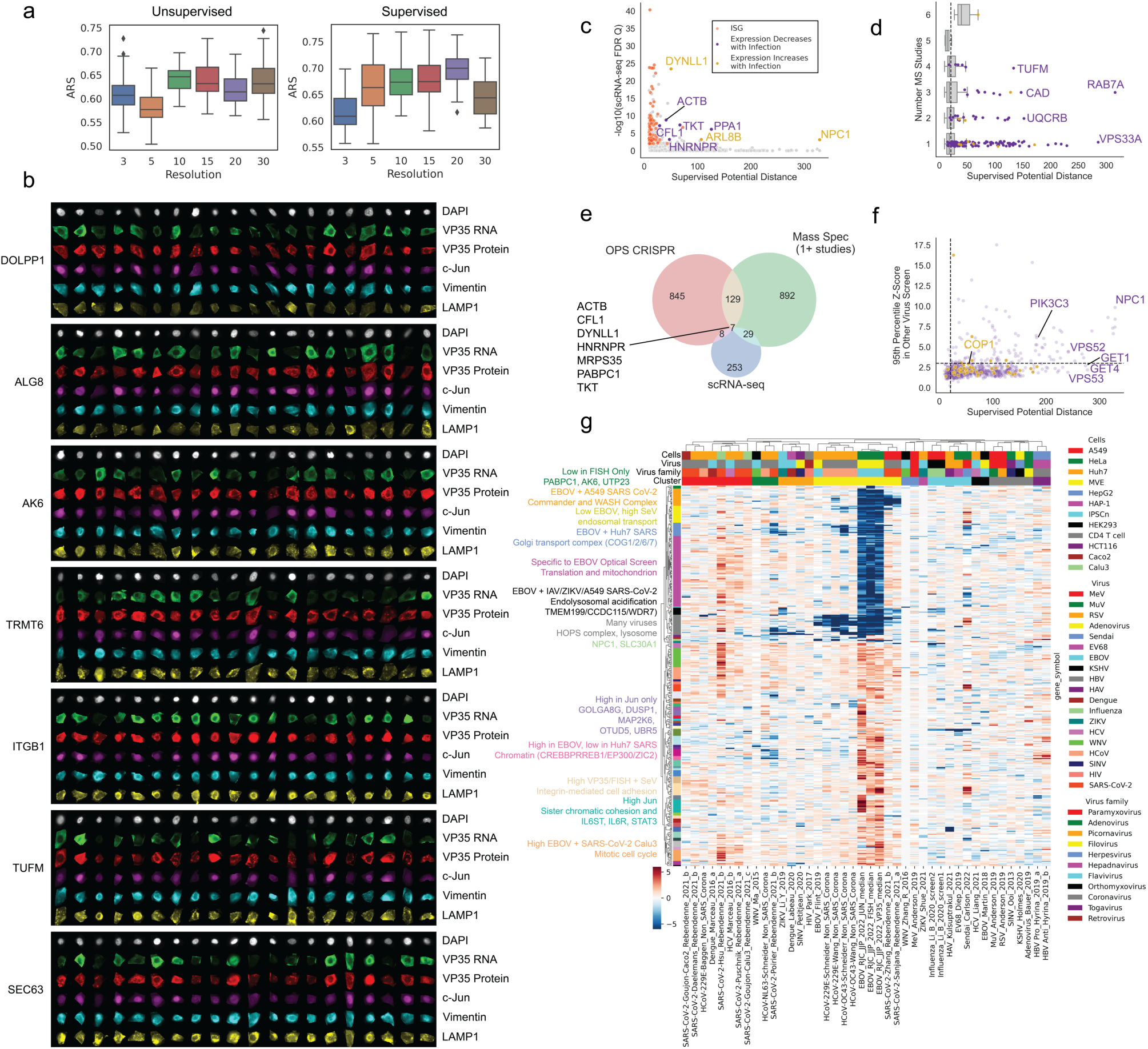
(A) Adjusted Rand score for Leiden clustering at different resolutions. (B) Additional single-cell images of select genetic knockouts from the genome-wide optical pooled screen. (C) Correlation between the PHATE potential distance from the clustering using the fine-tuned model and the adjusted FDR p-value from the Kotliar study, noting genes whose expression significantly increased or decreased along with infection. (D) Correlation between the PHATE potential distance from the supervised clustering and the number of mass spectrometry studies that identified the genes as an interactor with an Ebola virus protein. (E) Venn diagram showing overlap between top optical pooled screen hits, genes that were present in at least one mass spectrometry study, and differentially expressed genes from Kotliar et al’s single-cell RNA sequencing study. (F) Correlation between the PHATE potential distance from the supervised clustering and the 95th percentile z-score for each gene in other virus genetic screens, see (G). (G) Unsupervised clustering of hits from genome-wide virus genetic screens, hierarchical clustering performed using cosine distance.

**Figure S4.**
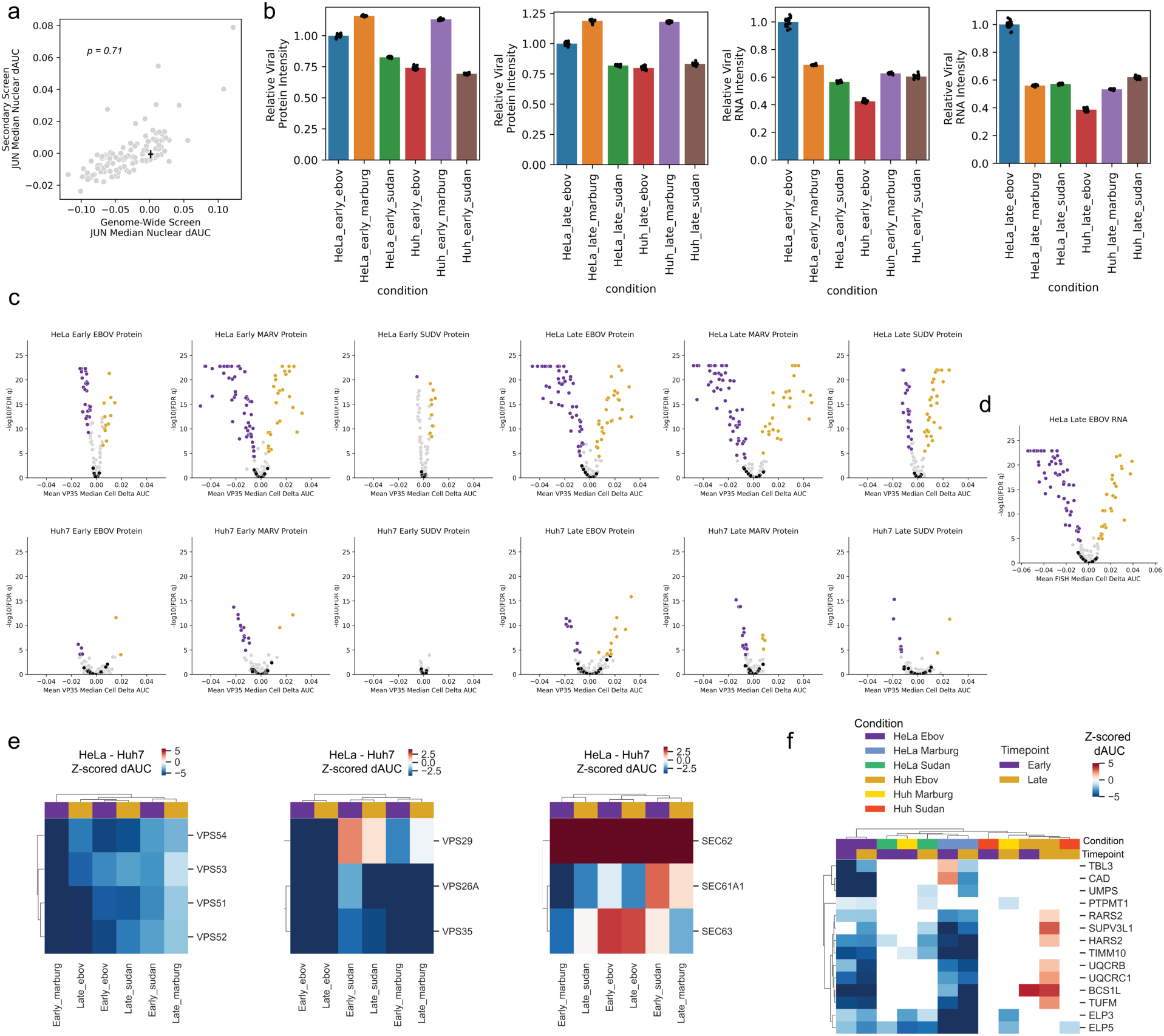
(A) Correlation between genome-wide c-Jun median nuclear delta AUC scores and secondary screen delta AUC scores; black lines indicate standard deviation for non-targeting control sgRNAs in each screen centered around the mean value for non-targeting sgRNAs in the screen. (B) Secondary screen mean viral protein (VP35 for EBOV and SUDV or VP40 for MARV) and RNA intensities in non-targeting control sgRNAs relative to HeLa cells infected with EBOV. (C) Volcano plots for VP35 (EBOV, SUDV) or VP40 (MARV) protein expression in each of the twelve screening conditions. (D) Volcano plot for viral VP35 RNA levels in HeLa cells at the late timepoint condition. (E) Heatmaps showing the difference between HeLa cell and Huh7 cell z-scored delta AUCs for members of the GARP, retromer, and the Sec61 complex. Hierarchical clustering performed using Euclidean distance. (F) Heatmap showing z-scored delta AUC values for genes identified as enriched for a punctate phenotype in the genome-wide screen and also included in secondary screens (white cells indicate conditions where p > 0.05). Hierarchical clustering performed using Pearson correlations.

